# Genomic evidence for phylogenetic congruence with sporadic host shifts in leishmaniaviruses and their parasite hosts

**DOI:** 10.1101/2025.11.20.685321

**Authors:** Senne Heeren, Lilian Motta Cantanhêde, Khaled Chourabi, Mayara Cristhine de Oliveira Santana, Joon Klaps, Alexei Kostygov, Vyacheslav Yurchenko, Philippe Lemey, Jean-Claude Dujardin, Frederik Van den Broeck, Elisa Cupolillo

**Affiliations:** Department of Biomedical Sciences, Institute of Tropical Medicine, Antwerp, Belgium; Department of Microbiology, Immunology and Transplantation, Rega Institute for Medical Research, Katholieke Universiteit Leuven, Leuven, Belgium; Department of Biomedical Sciences, University of Antwerp, Antwerp, Belgium; Leishmaniasis Research Laboratory, Oswaldo Cruz Institute, Oswaldo Cruz Foundation, Rio de Janeiro 21040360, Brazil; Instituto Nacional de Ciência e Tecnologia de Epidemiologia da Amazônia Ocidental, INCT EpiAmO, Porto Velho 76812100, Brazil; Life Science Research Centre, Faculty of Science, University of Ostrava, Ostrava, Czechia

**Keywords:** Co-evolution, *Leishmania*, *Leishmaniavirus*, phylogenetics

## Abstract

The association between *Leishmania* parasites and the double-stranded RNA virus, *Leishmaniavirus* (LRV), significantly impacts human health by modulating disease severity. Despite its clinical importance, the evolutionary history of this symbiosis remains largely unknown. This study presents one of the first formal investigations of their co-phylogenetic history across different taxonomic hierarchical levels; based on total RNA sequencing of cultivated isolates, simultaneously capturing the parasite’s transcriptome and the viral genome. We found strong support for phylogenetic congruence between *Leishmania* and LRV at both the subgenus and species level; based on genetic distance correlations and formal co-phylogenetic statistics. Focusing on the interactions of *L.* (*V.*) *braziliensis* and *L.* (*V.*) *guyanensis* with LRV1, we observed weaker co-phylogenetic signals though frequent instances of intraspecific host switching. Together, our findings indicate that LRV has been a persistent evolutionary partner of *Leishmania*, providing a framework for understanding the emergence and maintenance of virus-mediated disease phenotypes.

## Introduction

Interspecies interactions are key to ecosystem functioning. In this regard, viruses also play an important role in our biosphere, where they often cause direct harm to their hosts’ populations, sometimes leading to severe socio-economic crises on regional or global scales. Conversely, viruses may also affect populations indirectly through so-called nesting doll (“matryoshka”) infections ^1–6^. In such cases, viruses have established persistent, sometimes mutualistic-like, relationships with organisms such as pathogenic parasites, where they may alter (and often exacerbate) disease manifestation in humans. This may complicate treatment of the infection and ultimately hinder disease control efforts in endemic areas ^6–9^. While efforts are currently underway to assess the immediate clinical and epidemiological impact of such nesting-doll infections, these systems also serve as a powerful biological model for understanding the evolutionary resilience of endosymbiotic relationships. Investigating these “matryoshka” infections allow for determining whether a virus remains a transient passenger or becomes an integral part of the host’s biological identity; a process that fundamentally shapes how complex pathogens adapt and diversify over millions of years.

To address this, we investigated the shared evolutionary history of an iconic virus–parasite symbiotic system comprising human pathogenic *Leishmania* parasites (Trypanosomatidae), the causative agent of leishmaniasis, and their persistent double-stranded RNA viruses of the genus *Leishmaniavirus* (LRV) (*Pseudototiviridae, Duplornaviricota*) ^10,11^. In this system, LRV has been found to mutualistically aid the parasite by modulating the innate immune response of the mammalian host through an interaction with Toll-like receptor 3 ^7^ which, in turn, ensures the inhibition of the Nod-like receptor protein 3 inflammasome in host macrophages ^12^. This immunological cascade ultimately leads to increased parasite survival, disease progression, exacerbation, and an increased likelihood of treatment failure ^8,9,12,13^. In exchange, the virus’ own metabolism and replication are facilitated by several host-encoded proteins (e.g., kinases and proteases) ^14^. While most research on the impact of LRV on leishmaniasis infections has been conducted on LRV1 (see below), evidence of similar or differential effects caused by LRV2 on the disease severity, *Leishmania*’s virulence factors or overall phenotype is rather limited, though emerging ^15–18^.

Since the first trypanosomatid viruses (LRV1 and LRV2) were characterized on the genome level about 30-40 years ago, the *Leishmaniavirus* classification has expanded to encompass six species associated with either *Leishmania* or *Blechomonas* spp. ^1,11,19^. Within *Leishmania* specifically, three LRV species are currently described: *Leishmaniavirus ichi* (referred to below as “LRV1”) associated with New World *L.* (*Viannia*) spp., *Leishmaniavirus ni* (“LRV2”) associated with Old World *L.* (*Leishmania*) spp. and *L.* (*Sauroleishmania*) spp., and *Leishmaniavirus sani* associated with *L.* (*L.*) *aethiopica* (also Old World) ^2,11^. Despite numerous reports suggesting a long-standing co-divergence or sporadic interspecific host-switching, formal assessment of the co-evolutionary history, by means of co-phylogenetic inference methods, of *Leishmania* and *Leishmaniavirus* remains lacking ^1,10,20–22^. The recent population genomic study in Peru and Bolivia demonstrated that parasite gene flow and hybridization are key drivers of viral dissemination ^3^. In these regions, hybrid *Leishmania braziliensis* lineages were found to harbor a significantly higher prevalence and diversity of LRV1 compared to ancestral, reproductively isolated populations, suggesting that hybridization can “replenish” viral diversity and facilitate the spread of the virus into new ecological niches as well as hosts.

Here, we determined whether these sporadic host shifts are common enough to erode long-term phylogenetic congruence across different taxonomic levels. Because most historical studies focused on analyzing the virus or the parasite separately and relied on a limited number of genetic markers rather than taking a genome-wide approach, we present the most comprehensive co-phylogenetic assessment to date, across multiple hierarchical levels of taxonomy (subgenus, species, and population). We hypothesized that strong patterns of phylogenetic congruence (i.e., mirroring symbiont phylogenies) would remain intact at the higher subgeneric and species levels, while a more dynamic co-phylogenetic signal (i.e., having a tendency where closely related hosts are associated with closely related symbionts) would emerge at the population level. Resolving this is essential for understanding the evolutionary resilience of this symbiosis, ultimately providing a more general view on the evolutionary interplay between viruses and pathogenic protists.

## Results

### Expanding the known genomic diversity of LRV1 in South America

Starting from 159 *Leishmania* (*Viannia*) spp. isolates, PCR-screening revealed 66 LRV1-positive ones (Tab. 1; S. Fig. 1; S. Tab. 1). These were subjected to total RNA sequencing to obtain both the viral genomes and the parasite transcriptomes. Viral genomes were reconstructed based on a de novo assembly approach as implemented in a novel genome reconstruction pipeline for viral sequencing data, Viralmetagenome ^23^. With this pipeline, we were able to reconstruct viral genomes from 49 isolates, of which 42 were of sufficiently high quality, acceptable for subsequent co-phylogenetic analyses (S. Tab. 2). These 42 isolates originated from four Amazonian states of Brazil (Acre: n = 4; Amazonas: n = 21; Pará: n = 6; Rondônia: n = 10) and one Venezuelan state (Trujillo: n = 1), which were sampled between 1985 and 2017. The remaining seven isolates were not retained for subsequent analyses, as these reconstructed genomes were either too short in length (IOCL2398, IOCL3007, IOCL3515, IOCL3654) or harbored a higher number of SNPs when the sequencing reads were remapped to their respective assemblies (IOCL3562, IOCl3563, IOCL3567) (S. Tab. 2). For one isolate in particular, IOCL3567, the pipeline yielded three genomes of 4,667 – 5,185 bp with a median read depth of 370× – 372×, but with a high number of SNPs (over 600). This suggests infection by at least two LRV1 viruses, a phenomenon previously documented in other *Leishmania*-LRV1 systems ^3,24^. However, we were unable to further disentangle these chimeric-like assemblies and discarded this isolate from downstream analyses. The same was done for another 17 isolates, for which the assembly pipeline did not yield properly reconstructed LRV1 genomes.

**Tab. 1:** LRV1 prevalence across all 159 screened *Leishmania* (*Viannia*) spp.

| **Species** | **LRV1 prevalence** |
| --- | --- |
| *L. braziliensis* | 27.85% (22/79) |
| *L. guyanensis* | 75.61% (31/41) |
| *L. lainsoni* | 16.67% (1/6) |
| *L. lindenbergi* | 0% (0/6) |
| *L. naiffi* | 61.11% (11/18) |
| *L. panamensis* | 0% (0/4) |
| *L. shawi* | 33.33% (1/3) |
| *L. utingensis* | 0% (0/1) |
| *L. (Viannia) spp.* | 0% (0/1) |
| **Total** | **41.51% (66/159)** |

The 42 selected assemblies covered 67.6% to 99.6% of the LRV1-4 genome (5,284 bp; SRA NC_003601) serving as a reference (S. Tab. 2). The percentage of sequencing reads mapping back to the assemblies ranged from 0.0002% to 0.027% (median = 0.004%). The median depths of coverage for the assemblies were between 101× and 1,093×, with an overall median of 391× (S. Tab. 2). The analysis of variants revealed a median of 2 heterozygous SNPs (average = 8; min = 0; max = 84) and no homozygous SNPs in any of the samples, demonstrating the reliability of the reconstructed genomes. All 42 genomes resembled the typical arrangement of LRV1, with two overlapping open reading frames coding for a capsid protein (CP; 2,999 bp) and an RNA-dependent RNA polymerase (RDRP; 2,637 bp), which were near-completely covered in most cases (37 assemblies with a genome coverage over 95%; S. Tab. 2).

Phylogenetic reconstruction and calculation of pairwise genetic distances between the 42 assembled genomes revealed 14 highly supported viral clades (Fig. 1a) exhibiting relatively low, within-clade, genetic distances (<0.125 substitutions/site; S. Tab. 3). The clades were named according to their host species with Roman numerals (i.e., LgI-V, LbI-V, LsI, LnI-II) (Fig. 1). Analysis of geographic distribution revealed that, among the studied isolates, those from the Amazonas belonged to eight clades, three of which being unique to this location (LgIII, LnI, LsI) (Fig. 1b). Other clades found in a single state included LbII and LbIV in Acre; LbI and LbIII in Rondônia; and LgVI and LnII in Pará. The single LRV1 genome of Venezuelan origin belonged to the LgII-clade, which was mainly identified in Amazonas and Acre. However, the current sample size impeded our ability to infer the connection of *Leishmania* and LRV1 between Trujillo and the Western Brazilian states.

**Fig 1.**
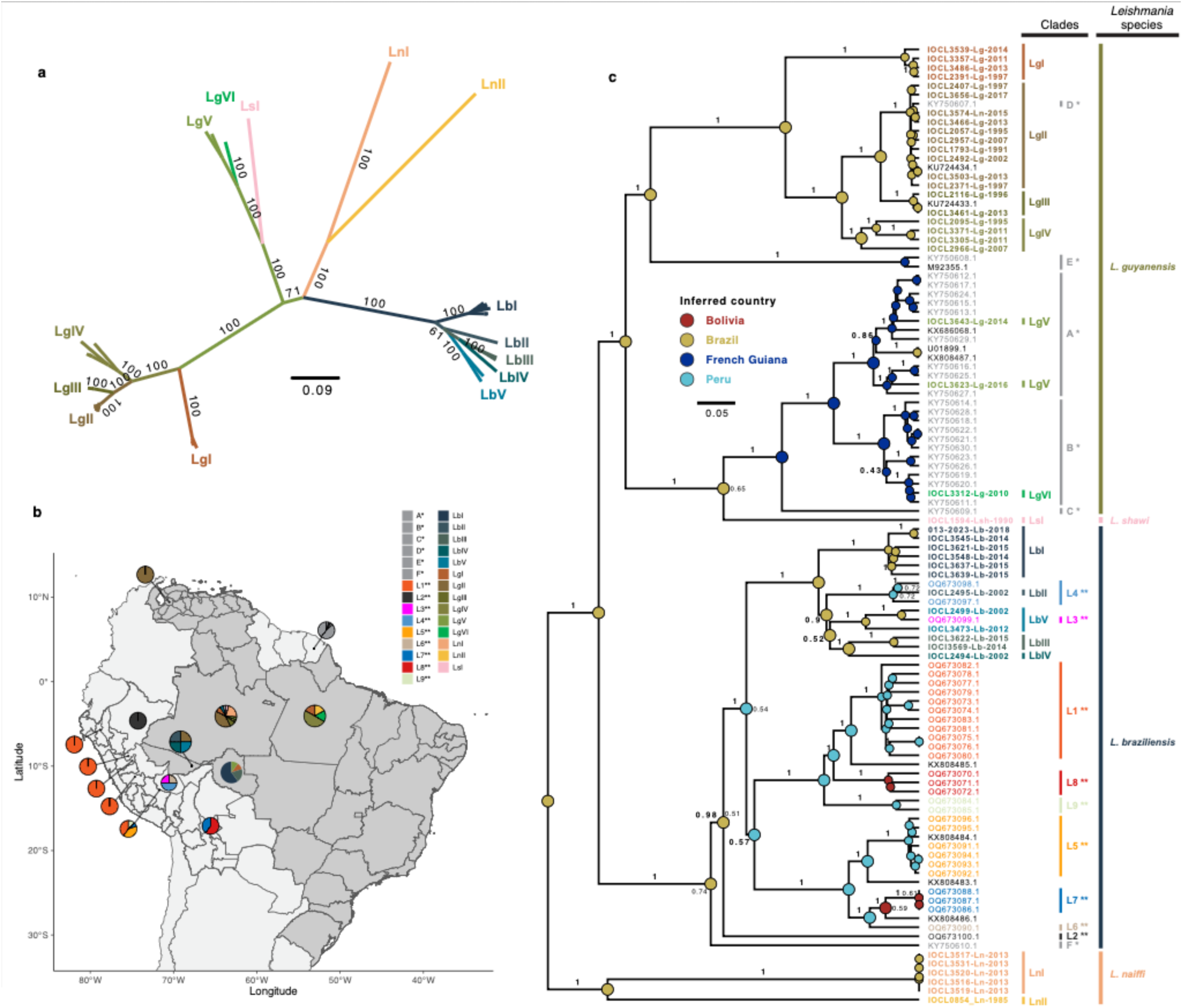
Lineage diversity and distribution of the LRV1 genomic diversity. **a** Unrooted maximum likelihood phylogenetic tree based on 42 near-whole genome LRV1 sequences. Branch values represent bootstrap support values based on 10,000 ultra-fast approximated bootstrap replicates. The scale bar depicts the number of substitutions per site. Clades were delineated based on high bootstrap support and low (within-clade) pairwise genetic distances (S. Tab. 3). Colors are set for visual purposes and do not serve any information on the ancestral states (i.e. internal nodes). The abbreviations Lb, Lg, Ln and Ls refer to the *Leishmania* hosts of the LRVs: *L. braziliensis*, *L. guyanensis*, *L. naiffi* and *L. shawi*, respectively. **b** Administrative map of South America showing the geographic origin of the 42 LRV1 genomes (dark gray area) along with geolocalized LRV1 genomes reported previously ^3,24^. Administrative country-and state-level data originate from: http://www.diva-gis.org/Data. **c** Midpoint rooted maximum clade credibility tree based on the 42 newly obtained near-whole LRV1 genomes along with 64 publicly available ones. Branch support values (bold faced) represent the posterior probabilities. Nodes are colored according to their inferred geographic origin, with posterior probabilities shown adjacently in regular font. Tip labels are colored according to the viral lineages (from this study or previous ones). Bars beside the tree represent the newly or previously described viral lineages (matching the tip labels) and the associated *Leishmania* species. Single and double asterisks mark the lineages identified by Tirera et al. (2017) ^24^ and Heeren et al. (2023) ^3^, respectively.

Our new panel of LRV1 genomes provides a major expansion of the known genomic diversity of the virus. This becomes clear from the phylogenetic reconstruction, combining our panel with 64 publicly available genomes. Most clades characterized above had not been identified earlier, whereas a few (LgII, LgV, LgVI, LbII, and LbV) clustered with previously described ones ^3,24^ (Fig. 1c). The concomitant inference of the ancestral geographic origins (Fig. 1c, internal nodes) along with the clear host specificity of LRV1 towards the different *Leishmania* species suggests spatial divergence and subsequent spread of each species-specific clade. One exception to this observed host-specificity is LRV1-Ln-IOCL3574, which clusters with viruses associated with *L. guyanensis* (Fig. 1c). The corresponding *Leishmania* isolate was originally typed as *L. naiffi*, but our whole transcriptome data indicated that it is a mixture containing genetic material of both *L. naiffi* and *L. guyanensis* (see below).

Phylogenetic reconstructions based on the single-gene alignments (i.e., CP or RDRP) were similar to the clade delimitations based on the whole-genome data (Fig. 1b; S. Fig. 2). Moreover, co-phylogenetic global-fit tests (Robinson-Foulds distance, ParaFit and PACo) revealed a consistently strong co-phylogenetic signal between the different LRV1 phylogenies (i.e., pairwise comparisons between the genome-wide, CP and RDRP alignments) (S. Tab. 5; S. Fig. 2). The only striking inconsistency was the internal position of *L. guyanensis*-associated LRV1 from French Guiana (clades: LgV, LgVI, LsI, A*, B*, D*, E* (except M92355.1, origin: Suriname)). Based on both the CP and genome-wide phylogenies, this clade (S. Fig. 2, clade shown in red) appears to share a common ancestor with the other *L. guyanensis*-associated LRV1 genomes, whereas it formed a monophyletic group with the *L. braziliensis*-associated LRV1 in the RDRP phylogeny. Nevertheless, this inconsistency does not refute the observed LRV1 host-specificity, it merely indicates the uncertainty of the clade divergence (i.e., low bootstrap supports) at these ancestral nodes based on the two single gene alignments (S. Fig. 2). In addition, none of the three alignments showed any indication of recombination, based on the pairwise homoplasy index test (whole genome: *p*-value = 1.0; CP: *p*-value = 0.95; RDRP: *p*-value = 1.0).

### Transcriptomic diversity of the *Leishmania* (*Viannia*) subgenus

Total RNA sequencing of the LRV1-positive *Leishmania* isolates allowed us not only to assemble LRV1 genomes, but also to genotype the *Leishmania* isolates themselves based on the transcriptomic data. Similar to our previous whole-genome sequencing studies ^25,26^, we compiled a dataset of 115 isolates containing (i) genomic data from 66 isolates, generated for this work and (ii) 49 publicly available sequences from various *Leishmania* spp. that served as reference strains during genotyping and species delimitation. This reference panel contained isolates from the *Leishmania* (*Viannia*) *braziliensis* clade (*Lb* L1: n = 18; *Lb* L2: n = 3; *Lb* L3: n = 2; *Lb* L4: n = 2; *L. peruviana*: n = 2), the *Leishmania* (*V.*) *guyanensis* species complex (*L. guyanensis*: n = 3; *L. panamensis*: n = 14; *L. shawi*: n = 1), *Leishmania* (*V*.) *naiffi* (n = 3), and *Leishmania* (*V*.) *lainsoni* (n = 1) (S. Tab. 6) ^25^.

As our data was derived from RNA sequencing, the genotyping analyses focused on the coding DNA sequence (CDS) regions. Mapping the reads against the long-read-based M2904 *L. braziliensis* reference genome ^27^ resulted in a median read depth of 17× (min = 1×; max = 151×) across the entire CDS for all 115 isolates. For the isolates from this study alone, the median CDS coverage was 10.5× (min = 1×; max = 43×). Considering a minimum CDS read depth of 10×, we excluded 31 isolates (30 from this study and one from a previous study) from downstream analyses and filtered out 1,184 individual CDS regions below this threshold in the retained samples. The final genotype dataset comprised 6,299 CDS regions across 84 isolates, yielding 157,649 high-quality SNPs.

Based on the differences in the number of SNPs (S. Tab. 8) between the included *Leishmania* (*Viannia*) spp. (Chi-squared = 76.92; df = 10; *p* = 2.01e-12; pairwise Dunn tests and effect sizes: S. Tab. 9), all isolates, with the exception of IOCL3539 and IOCL3574, could be identified (*L. braziliensis* L1: n = 12; *L. guyanensis*: n = 18; *L. naiffi*: n = 4) (Fig. 2a). The latter two isolates harbored distinctively higher numbers of heterozygous SNPs compared to all other isolates (Fig. 2a; S. Tab. 8) and showed reticulation in the CDS-wide phylogenetic network, a pattern suggestive of recombination and potential hybrid ancestry (Fig. 2b).

**Fig 2.**
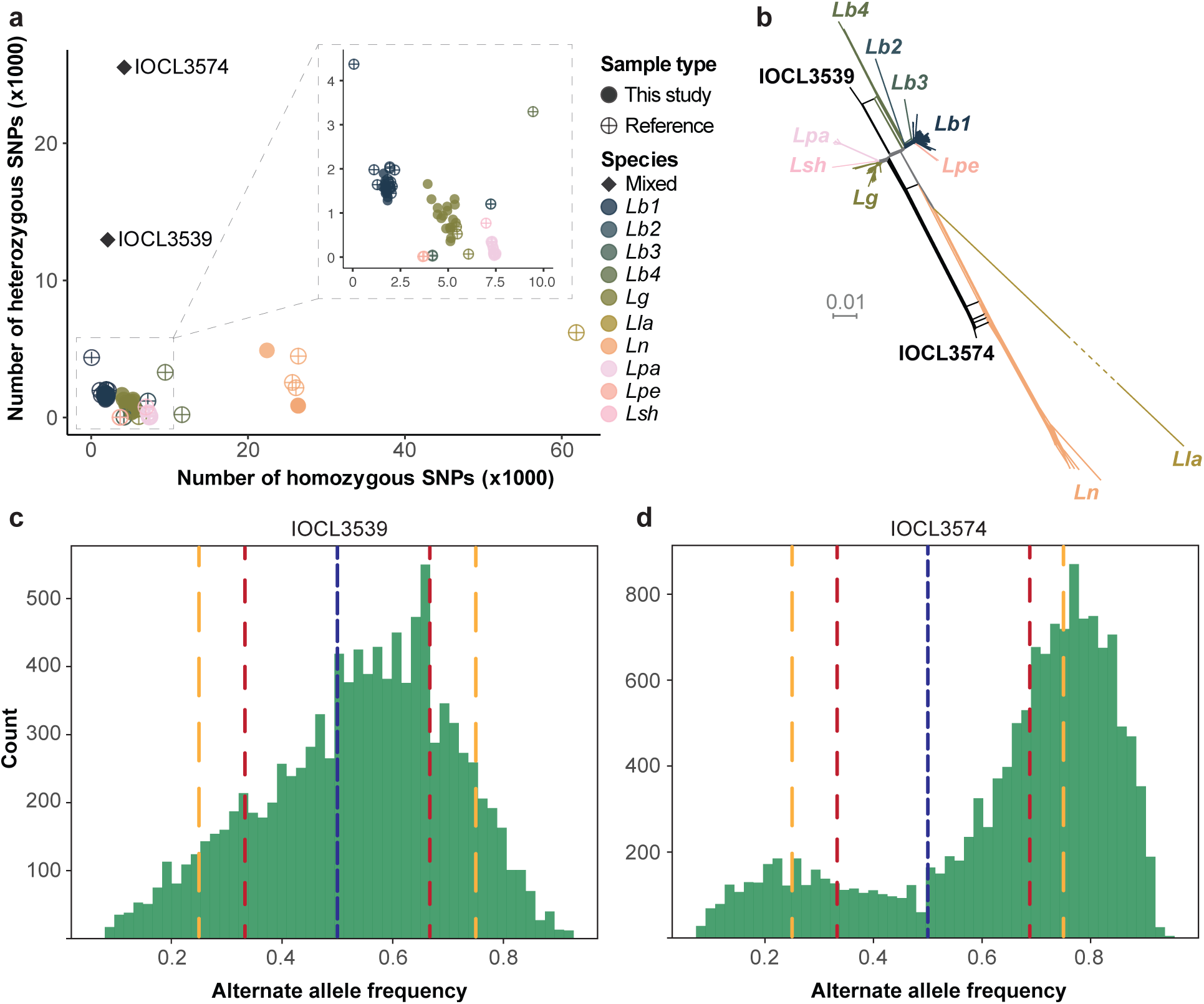
Species/lineage delimitation of 36 *Leishmania* isolates based on whole transcriptome data (157,649 bi-allelic SNPs). **a** Quantification of homozygous (x-axis) and heterozygous (y-axis) SNPs for 36 isolates in this study and 48 reference isolates (incl. the *L. braziliensis* clade, the *L. guyanensis* species complex, *L. naiffi* and *L. lainsoni*). **b** Phylogenetic network of all 84 isolates revealing the reticulate pattern caused by IOCL3539 and IOCL3574, indicative of hybrid ancestry. **c-d** Shifted CDS-wide alternate allele read depth frequency distribution plots of IOCL3539 (**c**) and IOCL3574 (**d**). Dashed vertical lines show the typical modalities for diploid (i.e., a unimodal distribution with a mode around 0.5; blue) and triploid (i.e., a bimodal distribution with modes around 0.33 and 0.66 in red) and tetraploid (i.e., the outer two modes around 0.25 and 0.75 in yellow, and the mode in-between at 0.5 in blue) genomes ^28^.

However, the alternate allele read depth frequency (ARDF) of these two isolates revealed uncommon profiles. In IOCL3539, the ARDF distribution was unimodal and skewed with a mode around 0.66, compared to the typical symmetric distribution around 0.5 ^28^ (Fig. 2c). The IOCL3574, on the other hand, showed a distribution that was bimodal with modes near 0.25 and 0.77. This contrasted with the typical bimodal ARDF distribution for triploid genomes (modes around 0.33 and 0.66) and was similar to the two outer modes that are typical for tetraploid genomes (0.25 and 0.75) but lacking its typical third mode at 0.5 (Fig. 2d). The CDS-wide shifts in the ARDF indicate that the observed hybrid-like patterns are the result of either the extensive mosaic aneuploidy, mixed infections or isolate contamination by two different *Leishmania* spp., rather than true interspecific hybrids ^29^. Moreover, the ARDF assessment on the chromosome-level revealed that both isolates were devoid of homozygous regions for the alternate allele, a pattern refuting the idea that these isolates are the result of true hybridization even more (S. Fig. 3). Because of the potentially artificial origin of the observed genomic pattern of these two isolates, both were excluded from the following downstream co-phylogenetic reconciliation analyses.

### Evidence for phylogenetic congruence across different hierarchical levels of *Leishmania* and LRV

To assess the extent to which *Leishmania* and LRV have shared a similar evolutionary trajectory, we performed a matrix regression on the pairwise genetic distances of both symbionts, combined with formal co-phylogenetic tests – global-fit and event-based – to detect the overall co-phylogenetic signal (i.e., the tendency of closely related hosts being associated with closely related symbionts) and evaluate true phylogenetic congruence (i.e., mirroring of symbiont phylogenies), respectively ^30^. These analyses were based on a dataset including 22 Peruvian *Leishmania* (*V.*) *braziliensis* isolates with their associated LRV1 genomes ^3^; as well as six Old World *Leishmania* isolates (*L.* (*L.*) *major* (n = 4), *L.* (*S.*) *tarentolae* (n = 1) and *L.* (*S.*) *adleri* (n = 1)) with their respective LRV2 genomes (S. Tab. 4,6).

On the subgeneric level, we observed a highly significant, strong, and positive correlation between pairwise genetic distances across the 51 *Leishmania*-LRV pairs that were analyzed (MMRR: R^2^ = 0.89; F = 10,913.49; *p* < 0.001) (Fig. 3a). In parallel, our array of global-fit analyses detected a consistently significant co-phylogenetic signal between *Leishmania* and LRV at this level (Fig. 3b; S. Tab. 10). Similarly, the event-based inference revealed strong signatures of phylogenetic congruence, suggesting that the two symbionts share a highly similar phylogenetic history (Fig. 3c). This latter analysis was conducted using nine different event-costs sets (S. Fig. 4a; S. Tab. 11), with eight of them yielding significant co-phylogenetic reconciliations, where well-supported co-speciation events (i.e., joint speciation/divergence of lineages that are ecologically associated ^31^) predominated the divergence of Old and New World *Leishmania* subgenera with LRV2 and LRV1, respectively (S. Fig. 5).

**Fig. 3.**
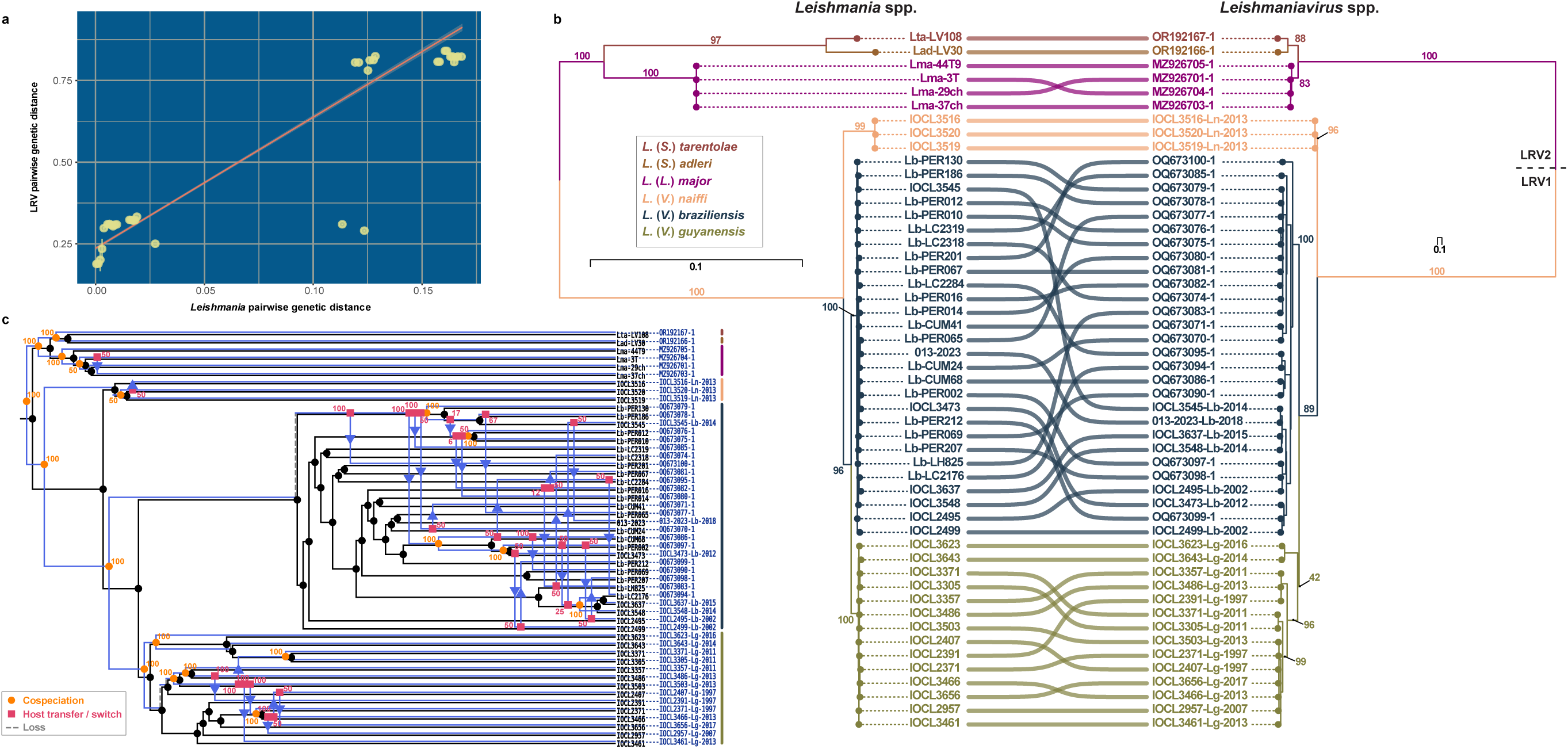
Co-phylogenetic reconciliation of *Leishmania*-LRV symbiosis at the subgeneric level. **a** Regression scatterplot of the pairwise genetic distances of *Leishmania* and LRV (51 pairs). Yellow points represent summarized values in 200 bins along the x-axis. **b** Tanglegram of *Leishmania* and LRV phylogenies. The *Leishmania* phylogeny was inferred from six concatenated gene sequences (S. Tab. 15; see methods). The LRV phylogeny was inferred from concatenated sequences of CP and RDRP (ORF2 and ORF3, respectively). Colors on the tree branches are set for visual purposes, indicative of the associated *Leishmania* species (tips), but do not serve any information on the ancestral states. Branch values represent bootstrap support values based on 10,000 ultra-fast approximated bootstrap replicates. The scale bar indicates the number of substitutions per site. **c** Event-based *Leishmania*-LRV co-phylogeny at the subgeneric level (51 tip-pairs). The reconciliation was inferred with the following event-costs: duplication = 1.50; transfer = 1.50; loss = 1.00 (S. Tab. 11; set 03). Support values for the inferred co-evolution events (e.g. co-speciation, host transfer) were estimated from 100 bootstrap replicates. Colored bars adjacent to the tip labels denote the LRV-host species as depicted in panel b.

When focusing on the species level (subgenus *L.* (*Viannia*)) and its association with LRV1, we observed a highly significant, yet more moderate, positive correlation between the parasite and viral pairwise genetic distances (MMRR: R^2^ = 0.59; F = 1,254; *p* < 0.001). Interestingly, the pronounced host specificity did not appear to be shaped by spatial separation, as viral genetic distances did not correlate with geographic distances between the isolates (MMRR: R^2^ = 0.005; F = 5.28; *p* = 0.135). In parallel, both the global-fit and event-based methods consistently detected strong patterns of co-phylogenetic signal and phylogenetic congruence, respectively (S. Tab. 10,12; S. Fig. 6). More specifically, the event-based analysis yielded a diverse set of possible co-phylogenetic histories based under six different event-cost scenarios (S. Fig. 4b; S. Tab. 12). Strikingly, in all cases, the inferred phylogenetic reconciliations were significant and, thus, supporting the presence of strong phylogenetic congruence between the members of the *Viannia* subgenus and LRV1 (S. Fig. 7). Another notable observation was that co-speciation events consistently accounted for species divergence (S. Tab. 12) and were well supported, whereas duplication (i.e., speciation of the symbiont lineage within one host lineage ^31^) and transfer events (i.e., host switch; transfer of symbiont from one host lineage to another ^31^) appeared to be more predominant within the separate species partitions (S. Fig. 7).

Finally, zooming on the population level of this virus-parasite co-evolutionary history, we focused on *L.* (*V.*) *braziliensis* and *L.* (*V.*) *guyanensis* with their respective LRV1 lineages. This also revealed signatures of phylogenetic congruence, albeit generally weaker, compared to the higher hierarchical levels (i.e., sub-genus and species levels), and occasionally lacking statistical significance. For instance, we found no significant correlation between the genetic distances of *L. braziliensis* and LRV1 (MMRR: R^2^ = 0.027; F = 10.65; *p* = 0.215) whilst a weak, though significant, correlation was observed in the case of *L. guyanensis* (MMRR: R^2^ = 0.349; F = 47.68; *p* = 0.002). Surprisingly, both global-fit and event-based analyses revealed significant co-phylogenetic signals and evidence of phylogenetic congruence between each *Leishmania* species and its associated LRV1 lineage (S. Fig. 4c-d, 8-10; S. Tab. 10, 13).

## Discussion

Our study provides the most extensive evaluation to date of the co-phylogenetic history of the *Leishmania*-LRV symbiosis, transforming our understanding of “matryoshka” infections from a clinical observation into a formal evolutionary model. While previous efforts relied on separate analyses of either the virus or the parasite, our joint evolutionary approach leveraged simultaneous genome-wide reconstructions for both symbionts. Although the genome-wide evidence is primarily derived from LRV1, the available LRV2 data support the same broad co-divergence framework at the subgenus level. Together, these findings support a model in which *Leishmaniavirus* appears to have predominantly co-diverged with its *Leishmania* hosts across evolutionary time, despite recurrent host-switching events at the population level as observed in *L.* (*V.*) *braziliensis* and *L.* (*V.*) *guyanensis*.

Before performing co-phylogenetic inferences, we conducted independent analyses on the genomic diversity of the newly sequenced *Leishmania* and LRV isolates. These preliminary steps allowed us to (i) complete isolate species typing and document host-specificity, (ii) identify potential hybrid and artificially mixed isolates and (iii) identify novel genomic variants and clades. Initial analyses identified a distinct clustering of *Leishmania* isolates; primarily *Leishmania (Viannia) braziliensis*, *L. (V.) guyanensis*, and *L. (V.) naiffi.* Two isolates (IOCL3539 and IOCL3574) showed patterns of inter-specific mixtures, either due to mixed infections or actual genomic hybridization. This is an important distinction as true hybridization (both inter-and intraspecific) is an increasingly recognized phenomenon in parasitic protists ^32,33^, and *Leishmania* in particular ^3,25–27,34–37^, with important epidemiological implications; consequently, the detection and reporting of new hybrids is essential within the context of genomic surveillance ^32,38^. Closer inspection, by means of evaluating the alternate allele read depth frequencies at heterozygous sites, revealed (i) shifts from the expected uni-, bi-, and trimodal patterns ^28^, and (ii) the absence of distinct homozygous regions for the alternate allele, contrasting with other previously reported hybrid signatures ^25,29^; genomic signatures that are indicative for mixed infections or extensive mosaic aneuploidy, rather than true hybrid signals. Owing to the tendency of these mixtures to be of “artificial” origin, both isolates were discarded from the downstream co-phylogenetic analyses.

Phylogenetic reconstruction of the newly assembled LRV1 genomes substantially expanded the known genomic diversity of the virus. This included (i) an increased intra-clade diversity, based on whole-genome data, within groups previously described ^3,24^, and (ii) the identification of novel clades (i.e., LgI, LgIII, LgIV, LbI, LbIII, LbIV, LsI, LnI, LnII). In concordance with Santana et al. (2023) ^39^, we observed strict host specificity among LRV1 genomes, independent of sampling location. This was supported by a strong correlation of virus genetic distances with those of their hosts, but not with geographic divergence at the species level within the *Leishmania* (*Viannia*) subgenus (with the sole exceptions being the presumably mixed isolates IOCL3539 and IOCL3574).

The main goal of this study was to formally validate (or refute) the long-standing hypotheses regarding the evolutionary association between *Leishmania* and LRV ^1,10,20–22^ within an explicit co-phylogenetic context, and with a particular focus on the New World (i.e., the *Leishmania* (*Viannia*) – LRV1 association). Overall, our findings revealed clear genomic evidence of virus-parasite phylogenetic congruence with strong support for predominant co-speciation events at both the subgenus (between the *Leishmania* and *Viannia* subgenera) and species (within the *Viannia* subgenus) levels. However, at the population level, this phylogenetic congruence was significantly reduced in both *L.* (*V.*) *braziliensis* and *L*. (*V*.) *guyanensis*. This shift coincides with our previous observations in Peru and Bolivia ^3^, where we found that parasite hybridization and gene flow act as key drivers of viral dissemination. In those regions, recombination between *Leishmania* lineages likely facilitates the’replenishing’ of viral diversity by allowing LRV1 to jump between previously isolated host populations. Our current results indicate that while these recombination-driven host shifts are common enough to weaken the co-phylogenetic signal at the population scale, they appear to remain sporadic on a macro-evolutionary level ^3,19,22,40–44^. It therefore seems that, despite frequent horizontal transfers, facilitated by the parasite’s exosomal pathway ^45–48^, vertical transmission remains the predominant factor shaping the long-term evolutionary trajectory of the *Leishmania*-LRV symbiosis ^2,10^, an interpretation that may be substantiated by taking into account the geographic population structure of the parasite hosts.

Multilocus microsatellite analyses previously revealed marked population structure among Brazilian *Leishmania* (*Viannia*), associated with both parasite species and geography, including distinct *L.* (*V.*) *braziliensis* populations from Amazonian and extra-Amazonian regions ^49^. More recently, genome-wide analyses confirmed pronounced geographic structuring within *L.* (*V.*) *braziliensis*, identifying two major parasite groups associated with the Amazon and Atlantic Forest biomes, in addition to smaller ecologically isolated populations ^26^. Although LRV1 occurrence was not investigated in these population genetic studies, this structure is noteworthy considering the known geographic distribution of the virus: LRV1 is frequently detected in Amazonian *Leishmania* (*Viannia*), whereas it appears to be absent or only sporadically detected in *L. braziliensis* from eastern and extra-Amazonian Brazil ^50^. Thus, the distribution of LRV1 broadly coincides with major components of the geographic population structure of its *Leishmania* hosts. Ecological differences among these regions may further reinforce this pattern. *Leishmania* (*Viannia*) transmission in Amazonia involves a diverse assemblage of sand fly vectors, whereas *L.* (*V.*) *braziliensis* transmission in Atlantic Forest and other extra-Amazonian settings involves distinct vector assemblages, with *Nyssomyia intermedia* playing an important role in several endemic areas of southeastern Brazil ^51^. Such geographic and ecological compartmentalization may restrict contact and gene flow among parasite populations and, consequently, reduce opportunities for horizontal LRV transfer across divergent *Leishmania* lineages. Therefore, the phylogenetic congruence observed here is consistent with long-term host-associated viral diversification and predominant co-divergence, but geographic host population structure and ecological filtering may also have contributed to generating and maintaining this pattern.

While our findings clearly indicate evidence for phylogenetic congruence between *Leishmania* and LRV, at least at the subgenus and species levels, this still does not provide proof for true virus-parasite codivergence or coevolution. Essentially, the presence of phylogenetic congruence patterns highlights the mirroring behaviour of the two symbiont phylogenies, which is essentially a topological feature disregarding individual divergence times as well as the temporal (in)consistency between the host-symbiont divergence ^30,52^. In other words, cophylogenetic reconciliation methods allow for the inference of temporally unrealistic events (i.e., “back-in-time” host switching) which can be explained by the computational complexity of implementing temporal constraints within these types of methods ^52,53^. This is a general limitation within phylogenetic reconciliation approaches which, consequently, makes it difficult to fully discern the presence of true codivergence from other alternative scenarios that could also result in patterns of (topological) phylogenetic congruence, such as pseudo-codivergence due to preferential host switching among more closely related hosts ^30,52,54,55^. Similarly, the observed decline in the overall co-phylogenetic signal of LRV1 at the population level of *L.* (*V.*) *braziliensis* and *L.* (*V.*) *guyanensis*, accompanied by the increased prevalence of inferred host switching events, does not provide unequivocal evidence that actual host switching has taken place. Alternatively, processes like host-range expansion (i.e., symbiont dispersal from one host lineage to another without loss in the ancestral lineage) ^55^, mixed infections of divergent viral lineages ^3,24^ or sorting events (in analogy to incomplete lineage sorting) ^56,57^ may give rise to similar patterns of lowered co-phylogenetic signal. Nevertheless, our findings do formally confirm the non-independence of the shared evolutionary history within this clinically important symbiosis indicating that LRV is a persistent evolutionary partner of *Leishmania,* providing a framework for understanding the emergence and maintenance of virus-mediated disease phenotypes.

It is also worth noting that this study does not elucidate the full co-evolutionary history of *Leishmania* and LRV. This is, in part, due to the discrepancy between the number of paired genomes originating from the Old-and New World regions. Although our analyses revealed strong signatures of phylogenetic congruence at the subgeneric level, these patterns warrant validation on a larger sample size of Old World isolates. Another unresolved question is whether LRV-presence is a plesio-(ancestral) or apomorphic (derived) trait in *Leishmania*. This is particularly relevant because of the absence of LRV in the divergent subgenus *Mundinia* (which harbours a phylogenetically divergent *Shilevirus* instead (*Leishbuviridae, Negarnaviricota*) ^58,59^). In other words, LRV could have been either lost in *L.* (*Mundinia*) or gained after its divergence from the most recent common ancestor of the other three subgenera (*Leishmania*, *Viannia*, and *Sauroleishmania*) ^10^. On top of that, elucidating the broader virus-parasite interplay is also becoming increasingly complex as more LRV species are described in the literature ^60^, such as the split of *Leishmania* RNA virus 2 in *Leishmaniavirus ni* and *L. sani* ^11^, and the LRV species that are associated with flea-infecting trypanosomatids of the genus *Blechomonas* ^19^. Moreover, LRV1 is not the sole RNA virus infecting *L.* (*Viannia*) spp. ^61^. Accounting for this growing symbiotic, co-evolutionary and taxonomic complexity, it is likely that additional, more cryptic, viral diversity remains undetected in these protistan parasites.

In conclusion, our total RNA sequencing approach enabled the simultaneous reconstruction of the evolutionary histories of *Leishmania* and their endosymbiotic LRV strains, with a primary focus on the *Leishmania Viannia* – LRV1 association. By simultaneously integrating parasite and viral phylogenies within a formal co-phylogenetic inference framework, we demonstrate that a robust signal of phylogenetic congruence – likely due to (pseudo)-cospeciation – persisted across millions of years of parasite evolution, spanning three different subgenera. This stability is punctuated by sporadic host shift-like events – mainly at the population level – likely facilitated by the parasite’s own sexual and parasexual dynamics. This work provides fundamental biological insight in the evolution of matryoshka infections where, ultimately, the host will pay the toll ^62^.

## Methods

### Parasite culturing & RNA sequencing

*Leishmania* isolates were cultured in vitro, screened for LRV1, and prepared for total RNA sequencing at the Fundação Oswaldo Cruz, Brazil (Fiocruz). Parasites were grown in liquid culture medium to a concentration of 10^7^–10^8^ cells/ml, collected through centrifugation (on the third day after the final subculturing step), and preserved in DNA/RNA Shield (Zymo Research). Total RNA was extracted using the PureLink RNA Mini Kit (Invitrogen), following the manufacturer’s instructions. All isolates were screened for the presence of LRV1 by PCR using distinct LRV1 primers: LRV F–5′-ATGCCTAAGAGTTTGGATTCG-3′ and LRV R–5′-ACAACCAGACGATTGCTGTG-3′ ^63^. The screening included a positive (*L. guyanensis* MHOM/BR/1975/M4147) and a negative control (*L. braziliensis* MHOM/BR/1975/M2903) ^64–66^. LRV1-positive isolates (n = 66) were sent for total RNA sequencing without polyA selection or rRNA depletion to make sure the LRV genome would not be lost in the sample, but could potentially affect the overall sequencing quality. Samples were sequenced on either the NovaSeq (n = 56) or DNBSeq (n = 10) platform at Genewiz (Azenta Life Sciences) and BGI Genomics, respectively. The latter ten samples were part of a pilot study to test the sequencing approach for capturing both the LRV genome and the parasite transcriptome. In both cases, paired-end sequences were generated to ∼30,000,000 150bp reads.

### *Leishmaniavirus* de novo assembly & phylogenetic reconstruction

Prior to the de novo assembly of viral genomes, raw reads from each of the 66 LRV-positive *L. Viannia* sequenced libraries were filtered and trimmed using fastp v.0.23.4 ^67^ (parameter settings: minimum base quality (-q): 30; percentage of unqualified bases (-u): 10; front-to-tail (-5) and tail-to-front (-3) per read sliding window trimming with window size (-W) of 1 and mean quality score (-M) of 30; right-cutting reads (--cut_right) per 10 bp windows (-- cut_right_window_size) when mean quality score (--cut_right_mean_quality) is below 30; only considering reads between 100 (-l) and 150 bp (-b)), and mapped against the *L. braziliensis* M2904 reference genome using SMALT v.0.7.6 (available at: https://www.sanger.ac.uk/tool/smalt-0/; parameter settings: k = 13; s = 2). Only unmapped reads were retained and processed with Viralmetagenome v.0.1.2 (available at: https://github.com/nf-core/viralmetagenome) for de novo reconstructions of viral genomes ^23^. In summary, with this workflow, we combined Megahit and Spades assemblies, followed by a BLAST search to extract contigs matching publicly available LRV1 genomes ^68–70^. Next, all “raw” assemblies were iteratively refined by re-mapping reads to the assembled genomes and subsequent variant and consensus calling, making use of BCFtools and iVar ^71,72^. The reconstruction of hybrid consensus genomes was skipped (--skip_hybrid_consensus: true). A detailed overview of the non-default parameter settings that were used is provided in the supplementary material (S. Tab. 14). Viralmetagenome was able to resolve a consensus genome for 49 of the 66 LRV1-positive isolates. Additionally, we conducted a post-hoc assessment of the assembly quality which involved re-mapping of the reads against their respective assemblies, and variant calling. In total, 42 assemblies were retained based on the following criteria: length > 3,000bp, read depth > 100×, and low SNP counts (i.e., <100 heterozygous and zero homozygous SNPs). Finally, whole-genome sequences, as well as sequences for the Capsid Protein (CP) and the RNA-dependent RNA Polymerase (RDRP), were aligned using the L-INS-I algorithm, as implemented in MAFFT v.7.490 (‘--localpair’ option) ^73^.

Phylogenetic reconstructions of LRV1 were generated using IQtree v.2.3.6 ^74^ over 10,000 bootstrap replicates using the ultrafast bootstrap approximation method ^75^. Substitution models were automatically selected by IQtree’s ModelFinder algorithm (-m MFP) ^76^, which selects the best substitution model based on the Bayesian Information Criterion after evaluating all models that are implemented in IQtree. Whole genome maximum likelihood phylogenies of the assembled LRV1 genomes were generated with and without a set of 64 publicly available genomes (S. Tab. 4) using GTR+F+I+R4 and GTR+F+I+G4, as respective substitution models. Similarly, gene specific phylogenies, including the 64 publicly available sequences, were inferred using the GTR+F+I+R4 substitution model for both CP and RDRP. For the clade delimitation of the 42 newly assembled genomes, we also calculated the pairwise genetic distances using the’dist.dna’ function of the R-package ape v.5.8-1 ^77^ (model =’raw’). Finally, recombination was tested by means of PHI tests (Pairwise Homoplasy Index), as implemented in SplitsTree v.4.17.0 ^78,79^.

In addition to regular phylogenetic reconstructions, we also attempted to reconstruct ancestral states across the entire LRV1 phylogeny, focusing on the virus’ geographic origin and associated *Leishmania* species. This was achieved by adopting a Bayesian phylogenetic framework using the BEAST v.1.10.4 software package ^80^ and the BEAGLE likelihood calculation library ^81^. Consistent with previous analyses ^3^, no temporal signal was detected in the data. Therefore, all sampling dates were set to zero, after which LRV1’s discrete phylogeography was inferred without time calibration, using the default strict molecular clock model. The nucleotide substitution model was set to GTR with estimated base frequencies and the site heterogeneity model set to a Gamma distribution with 4 discrete rate categories and accounting for invariable sites ^82–84^. The tree prior model was set to a speciation model, the Yule process ^85^, starting the analysis with a random starting tree (default setting). Two discrete traits were added: the country of origin and associated *Leishmania* species. For both traits, we selected the symmetric discrete trait substitution model along with the BSSVS (Bayesian Stochastic Search Variable Selection procedure) option in order to reconstruct their ancestral states. Prior and operator settings were kept at their default settings. The MCMC chain was run for ten million states, sampled every 1,000 states and providing a 10% burn-in. Throughout the inference process, convergence of parameter estimates was evaluated using BEAST’s auxiliary program Tracer v.1.7.2 ^86^. Upon completion of the runs, all effective sample sizes (ESS) of all parameters exceeded the recommended threshold of 200, except for the C-T transition rate in the GTR model, which reached an ESS of 110. Finally, the posterior tree distribution was summarized as a maximum clade credibility (MCC) tree using BEAST’s TreeAnnotator.

### Transcriptomic genotyping & phylogenetic reconstruction of *Leishmania* (*Viannia*) isolates

Total RNA sequencing reads of the 66 LRV1-positive *Leishmania* (*Viannia*) isolates were mapped against the *L. braziliensis* M2904 (MHOM/BR/75/M2904) reference genome (available at: https://tritrypdb.org/common/downloads/release-65/LbraziliensisMHOMBR75M2904_2019) using SMALT v.0.7.6 (same parameters as described above). Reads with a mapping quality lower than 25 were removed using SAMtools v.1.21 (‘-q’ option) and read depths across all positions were extracted (‘samtools depth-a’) ^71^. This reference genome has been shown to be useful for genotyping not only *L. braziliensis* isolates, but also other *L.* (*Viannia*) species, resulting in highly accessible genomes for downstream analyses ^25–27^. Because transcriptomic, rather than genomic, data were used, we discarded isolates with low median coverages (<10×) across the coding DNA sequences (CDS) from downstream analyses. Additionally, all CDS regions in every sample, with a coverage below 10× were omitted, ensuring that only transcriptomic regions with sufficient depth were included in the subsequent analyses. Variant calling (i.e., SNPs and INDELs) was carried out using GATK v.4.1.4.1 ^87^. Here, the initial calling was performed on a per-sample basis using HaplotypeCaller, after which all individual genotype vcf files were combined using CombineGVCFs, and variants were jointly genotyped with GenotypeGVCFs. Next, SNPs and INDELs were separated using SelectVariants, after which both panels were subjected to hard filtering with VariantFiltration. For SNPs, the following filters were applied: (i) QD < 2.0, FS > 60.0, MQ < 40.0, MQRankSum < −12.5 and ReadPosRankSum < −8.0, following GATK’s recommendations ^88^.

A phylogenetic network was generated using SplitsTree ^78^ based on the pairwise uncorrected *p*-distances calculated across 157,649 bi-allelic SNPs. This network was built using the NeighborNet ^89^ and EqualAngle ^90^ algorithms, as implemented in SplitsTree. Forty-nine publicly available isolates were included as reference sequences: members of the *L.* (*V.*) *braziliensis* clade, the *L.* (*V.*) *guyanensis* species complex, as well as *L.* (*V.*) *naiffi* and *L.* (*V.*) *lainsoni* (S. Tab. 6). In addition, the alternate allele read depth frequency (ARDF) distributions at heterozygous sites were calculated and visualized to identify whether potential hybrid isolates were the result from a mixed infection. This was done using a custom python script (vcf2freq.py; available at: https://github.com/FreBio/mytools). Differences in SNP counts between the different species were compared using a non-parametric Kruskal-Wallis (stats R-package ^91^) test, followed by (i) pairwise Dunn’s tests with additional Benjamini-Hochberg *p*-value correction for multiple testing (FSA R-package ^92^), and pairwise calculation of effect sizes (hedge’s g and the overall median difference), with 95% confidence intervals, as implemented in the dabestr R-package ^93^.

### Preparing alignments and phylogenies for co-phylogenetic reconciliation

Phylogenetic reconciliations of the LRV-*Leishmania* symbiosis were performed at the subgenus (combining Old World and New World species), species (focusing on the South American *Leishmania* (*Viannia*) species), population (focusing on *L.* (*V.*) *braziliensis* and *L.* (*V.*) *guyanensis* with their respective LRV1 lineages), and gene (reconciling the CP and RDRP gene phylogenies within LRV1) levels.

For the subgenus level analysis, we included 22 publicly available Peruvian *L.* (*V.*) *braziliensis* isolates and six Old World *Leishmania* isolates positive for LRV2 (S. Tab. 4,6). Instead of using the entire genome sequence, we concatenated the annotated CP and RDRP genes (i.e., ORFs 2 and 3 in the LRV genome) and aligned their sequences from LRV1 and LRV2 using the MAFFT v.7.490 ^73^ as described above. The LRV phylogeny was reconstructed using IQtree with 10,000 ultra-fast bootstrap replicates and the GTR+F+I+R3 substitution model based on the ModelFinder algorithm ^74–76^. For the maximum likelihood phylogenetic reconstruction of the *Leishmania* phylogeny, which included highly divergent species from different subgenera, a concatenated alignment of six orthologous protein coding genes was used (S. Tab. 15), as in previous publication ^94^. The sequences of the six target genes were obtained as follows. Reads were trimmed with fastp ^67^, using the same settings as indicated above, after which contigs were assembled (de novo) using Megahit v.1.2.9 ^68^. Contigs with six target genes were identified using a blastn ^70^ search and subsequently extracted. These sequences were combined with 222 publicly available ones (accession numbers listed in S. Tab. 15) and aligned using MAFFT, as described above. Finally, a phylogenetic tree was inferred using IQtree with 10,000 ultra-fast bootstrap replicates and TIM2+F+G4 set as optimal substitution model ^74–76^.

For the reconstruction of the parasite phylogenies at the species-and population-level co-phylogenetic analyses, we used concatenated SNP-alignments. Variants were called against the MHOM/BR/75/M2904 reference genome, as described above, resulting in genome-wide alignment containing 97,530 bi-allelic SNPs. From this panel, a phylogenetic tree was inferred using IQtree with 10,000 ultra-fast bootstrap replicates and the TVM+F+R2 as the most optimal substitution model, correcting for ascertainment bias (+ASC) ^74–76^. For the population-level analyses of *L. braziliensis*–LRV1 and *L. guyanensis*–LRV1, species specific partitions of the inferred *L.* (*Viannia*) phylogeny were used. Finally, for the reconciliation of the LRV1 CP and RDRP genes, we made use of the reconstructed phylogenies as described above.

### Genetic distance regression

In order to obtain an initial assessment of similarity between the viral and parasite phylogenies, we calculated pairwise genetic distances from the corresponding concatenated sequences using the ‘dist.dna’ function of the ape R-package v.5.0 ^77^ (model = ‘K80’). These distances were converted into distance matrices and analyzed using Multiple Matrix Regression with Randomization (MMRR) ^95^ where *p*-values were estimated over 1,000 permutations.

### Global-fit co-phylogenetic analyses

The presence of co-phylogenetic signals (i.e., testing whether closely related hosts tend to associate with closely related symbionts) between the parasite and virus phylogenies by making use of three global-fit methods was tested via (i) the Robinson-Foulds distance as implemented in the phytools R-package v.2.4-4 ^96^; (ii) ParaFit as implemented in ape ^77^, and (iii) PACo as implemented in the paco R-package v.0.4.2 ^97^. For each method, both virus and parasite phylogenies were downsampled using the ‘drop.tip’ function of ape. This way, only the isolates, for which both high-quality LRV1 genomes and *L.* (*Viannia*) transcriptome-based genotypes were available. Each approach was performed by running 1,000 permutations for *p*-value estimation and the Cailliez correction method was used in ParaFit and PACo to correct for negative eigenvalues.

### Event-based co-phylogenetic analysis

To test whether *Leishmania* and LRV exhibit signs of phylogenetic congruence, at one or multiple hierarchical levels, we made use of eMPRess, an event-based phylogenetic reconciliation tool used for reconciling phylogenies based on a set of events such as co-speciation, loss, duplication, and host-switch ^30,53–55^. A key advantage of eMPRess over other event-based reconciliation tools is its ability to specify different sets of event-costs prior to calculating the reconciliation, allowing for the evaluation of alternative evolutionary scenarios for obtaining more robust inferences ^53^. This is important as different event-costs schemes can lead to distinct solutions and interpretations. For each level, we visualized the cost-space (S. Fig. 4) and selected multiple sets of event-costs covering most of the cost-space (S. Tab. 11-13). Phylogenetic reconciliations were then calculated for each set, and a median MPR (i.e., a median value of the MPR solution space; MPR = Maximum Parsimony reconciliation) was visualized. Significance was assessed using 1,000 random permutations. Visualizations of the cost-spaces and reconciled phylogenies were generated using the eMPRess GUI, while significance testing was conducted with the CLI version (‘*p*-value’ function with ‘--n-samples 1000’ option).

## Acknowledgements

This work received financial support from the Directie-Generaal Ontwikkelingssamenwerking en Humanitaire Hulp (DGD) (Belgian cooperation). F.V.d.B. and S.H. acknowledge support from the Research Foundation Flanders (grants 1226120N and G092921N). S.H. additionally acknowledges support from KU Leuven’s Internal Funds (PDMt1/25/007). E.C. acknowledges support from Fundação Carlos Chagas Filho de Amparo à Pesquisa do Estado do Rio de Janeiro: CNE - 204.165/2024; 269.891/2021; ColBio - 210.285/2021; Conselho Nacional de Desenvolvimento Científico e Tecnológico: Research Fellow - 309627/2021-4; PROEP/IOC - 441639/2024-0; INCT-EpiAmo: 465657/2014-1; and the GGBN award: GGI-GGBN-2021-278.

L.M.C. acknowledges support from Fundação Carlos Chagas Filho de Amparo à Pesquisa do Estado do Rio de Janeiro, Pós-Doutorado; Nota 10 E-26/205.730/2022 and 205.731/2022. E.C., L.M.C, M.C.d.O.S. and K.C. also acknowledge support from the Coordenação de Aperfeiçoamento de Pessoal de Nível Superior – Brasil (CAPES) – Finance Code 001. V.Y. and A.Y.K. acknowledge support from the European Union through the Operational Programme “Just Transition” and the Czech Ministry of Environment (CZ.10.03.01/00/22_003/0000003 LERCO), as well as the Czech Science Foundation (grant 24-10009S).

## Data Availability

Sequencing data that were generated for this study, or not yet publicly available, were deposited on NCBI’s SRA under BioProject no. PRJNA1492813. The reconstructed LRV1 genomes, based on the same sequencing data, are deposited on NCBI’s GenBank under the following accession numbers: PZ723939 - PZ723980. Accession numbers of publicly available data that was used in complement are disclosed in the supplementary data (S.Table 1, 4, 6, 15).

## Code Availability

Scripts and data related to the analyses conducted in the light of this work are made publicly available on both Github (https://github.com/sheerenbiol/LRV_Leish_PhyloCon).

## Author Contributions

**Conceptualization**: S.H., E.C., F.V.D.B., J.C.D.; **Data curation**: S.H., K.C.; **Methodology**: S.H., L.M.C., K.C., M.C.d.O.S.; **Resources**: L.M.C., K.C., M.C.d.O.S., J.K., A.K., V.Y., E.C.; **Formal analysis**: S.H.; **Supervision:** J.C.D., E.C., P.L., F.V.D.B.; **Writing - original draft**: S.H., F.V.D.B., E.C.; **Writing - review & editing**: S.H., L.M.C, J.K., A.K., V.Y., P.L., J.C.D., F.V.D.B., E.C.

## Supplemental Information (SI)

**S. Fig. 1.**
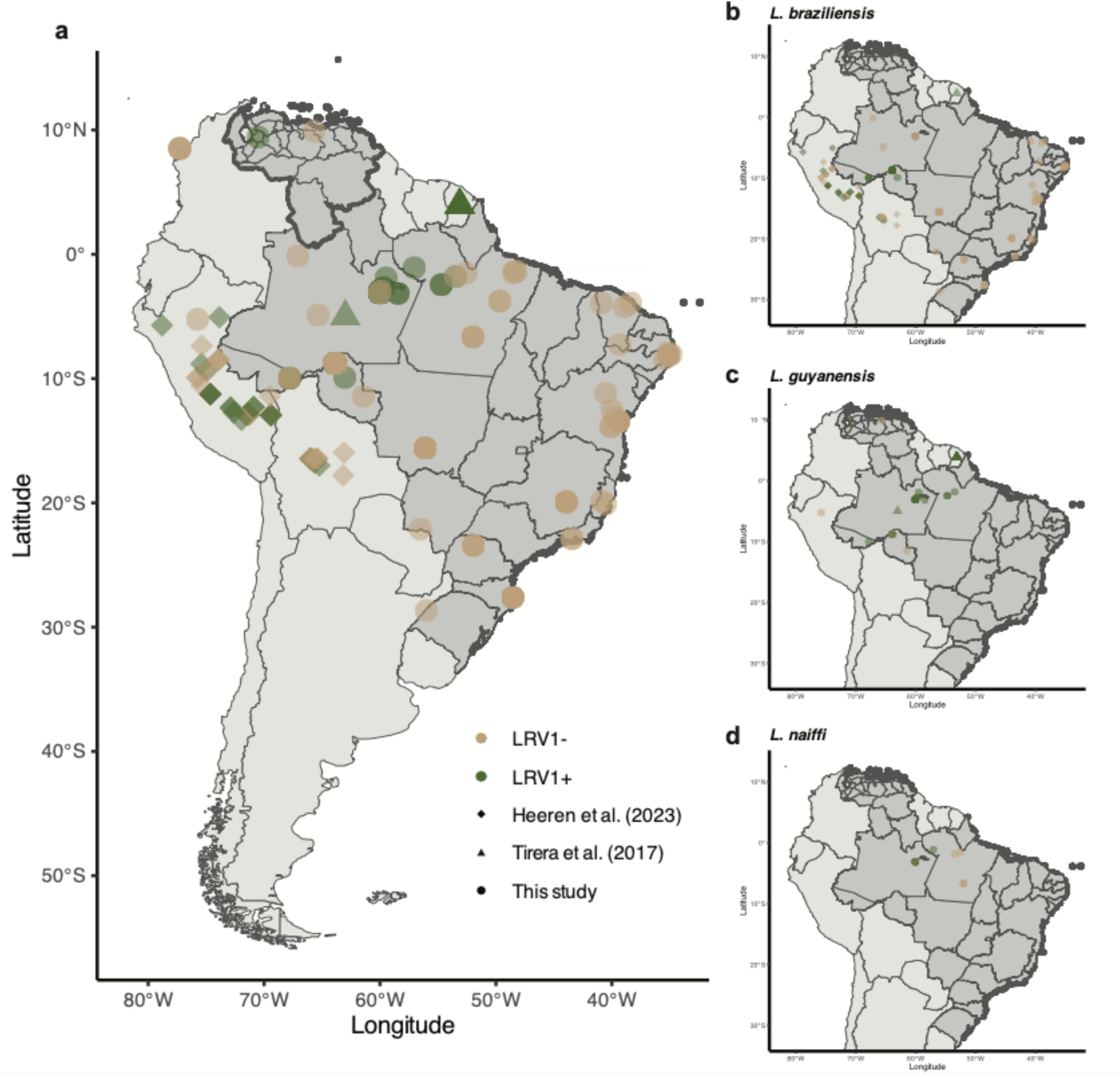
Spatial distribution of LRV1-positive and negative *Leishmania* isolates. **a** Spatial distribution of all isolates used in this study along with geolocalized *Leishmania* isolates from Heeren et al. (2023) ^3^ and Tirera et al. (2017) ^24^. **b-d** Spatial distribution of specific *Leishmania* species: *L. braziliensis* (**b**), *L. guyanensis* (**c**) and *L. naiffi* (**d**). Administrative country-and state-level data originate from: http://www.diva-gis.org/Data.

**S. Fig. 2.**
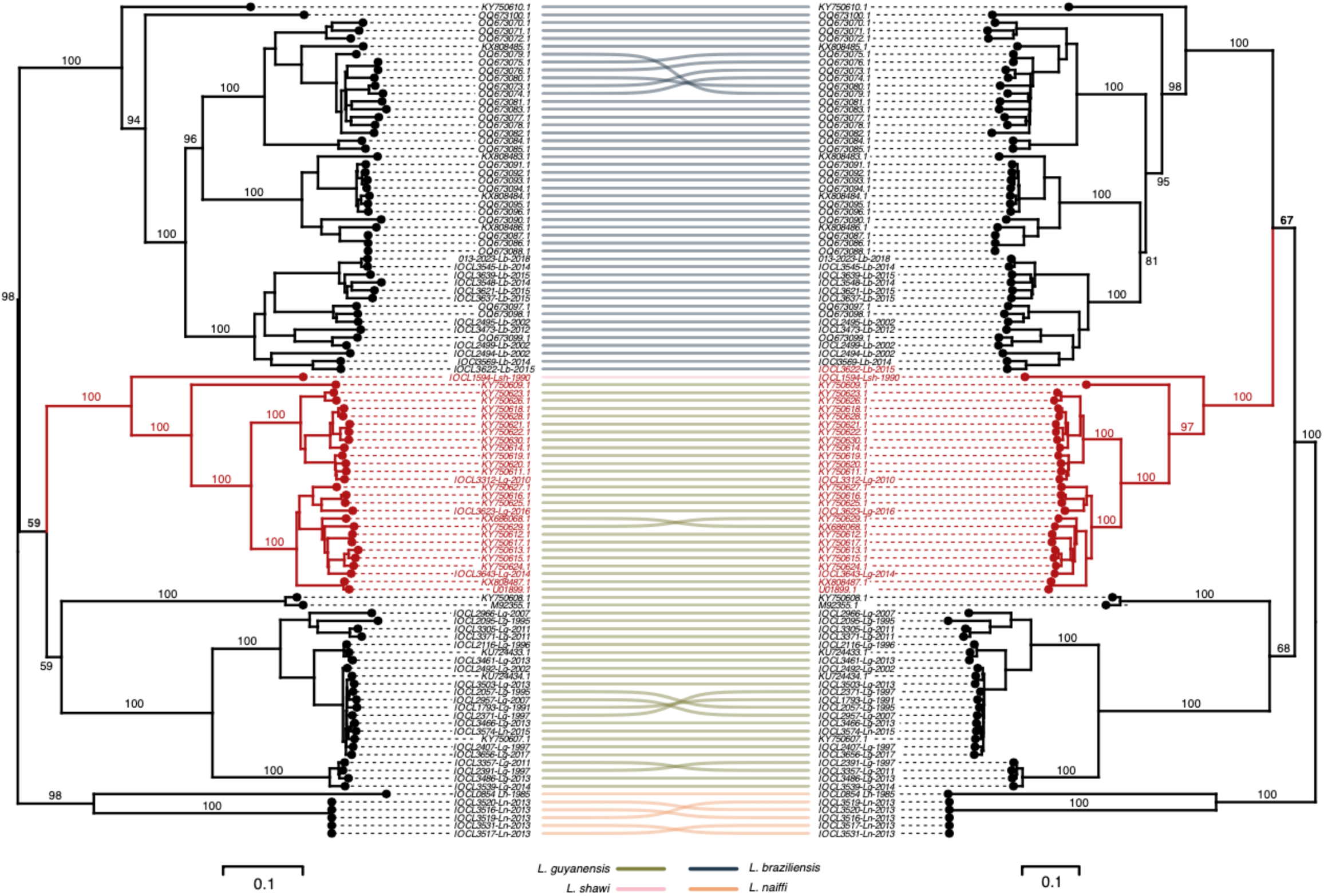
Tanglegram of the LRV1 capsid protein (CP) and RNA-dependent RNA polymerase (RDRP) gene phylogenies. The clade colored in red occupies different positions in the CP and RDRP phylogenies. Branch values represent bootstrap support values based on 10,000 ultra-fast approximated bootstrap replicates. The scale bar depicts the number of substitutions per site. Links between the trees are colored according to the identified clades of the 42 newly assembled LRV1 genomes (Figure 1; Suppl Figure 1).

**S. Fig. 3.**
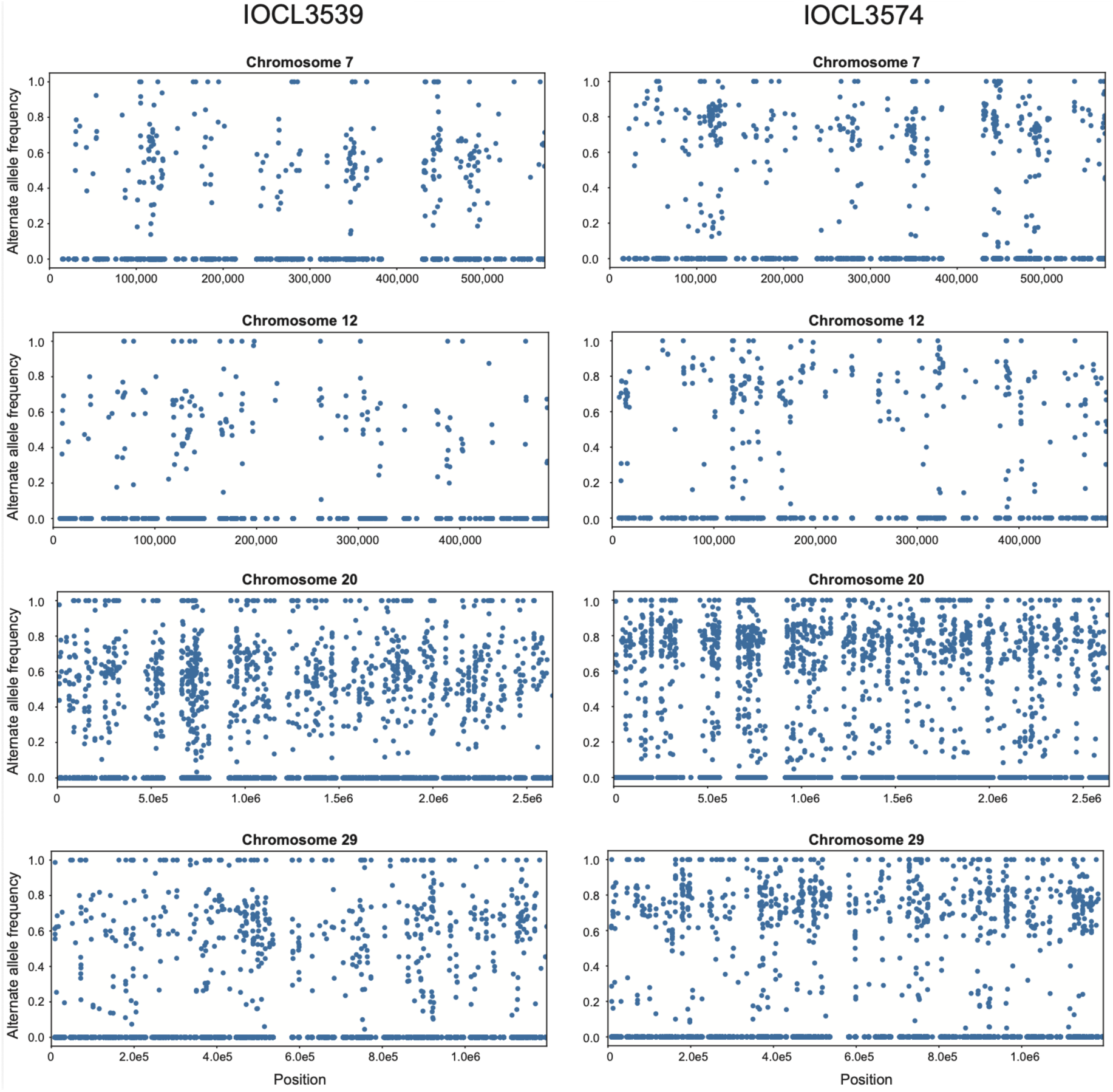
Chromosome-level read depth frequency distribution plots for the alternate allele for IOCL3539 (**left**) and IOCL3574 (**right**). X-axis indicates the position of each along the chromosome, while the y-axis denotes the read depth frequency of the alternate allele that is present in that SNP. Only chromosomes 7, 12, 20 and 29 are shown to exemplify the heterozygous nature of both isolates which are both devoid of homozygous stretches across their genomes.

**S. Fig. 4.**
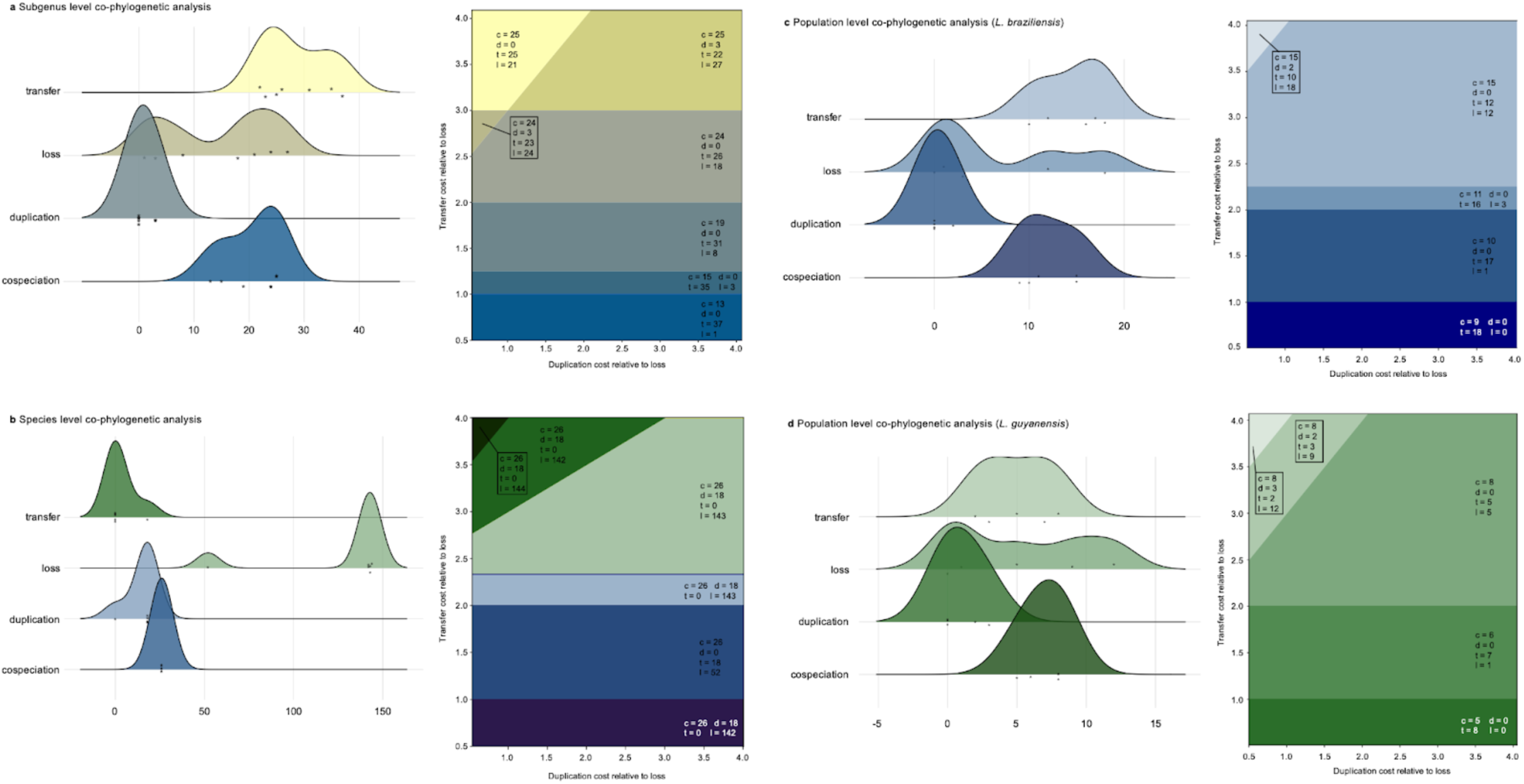
Event-cost distributions and cost spaces from event-based co-phylogenetic analyses of the *Leishmania*–LRV symbiosis across the different hierarchical levels. Per panel, the left-side plot depicts the density of the number of events that have been calculated over all the tested cost-sets; (ii) the right-side plot indicates the costspace as calculated by eMPRess. Each colored zone corresponds to the same number of events and reconciliation solutions. **a** Subgenus level. **b** Species level (within *L.* (*Viannia*)). Note the high number of inferred loss events in this analysis, resulting from an adjustment of the default loss cost from 1 to 0 to enable co-phylogenetic reconciliations in eMPRess. **c** Population level for *L.* (*V.*) *braziliensis*. **d** Population level for *L.* (*V.*) *guyanensis*.

**S. Fig. 5.**
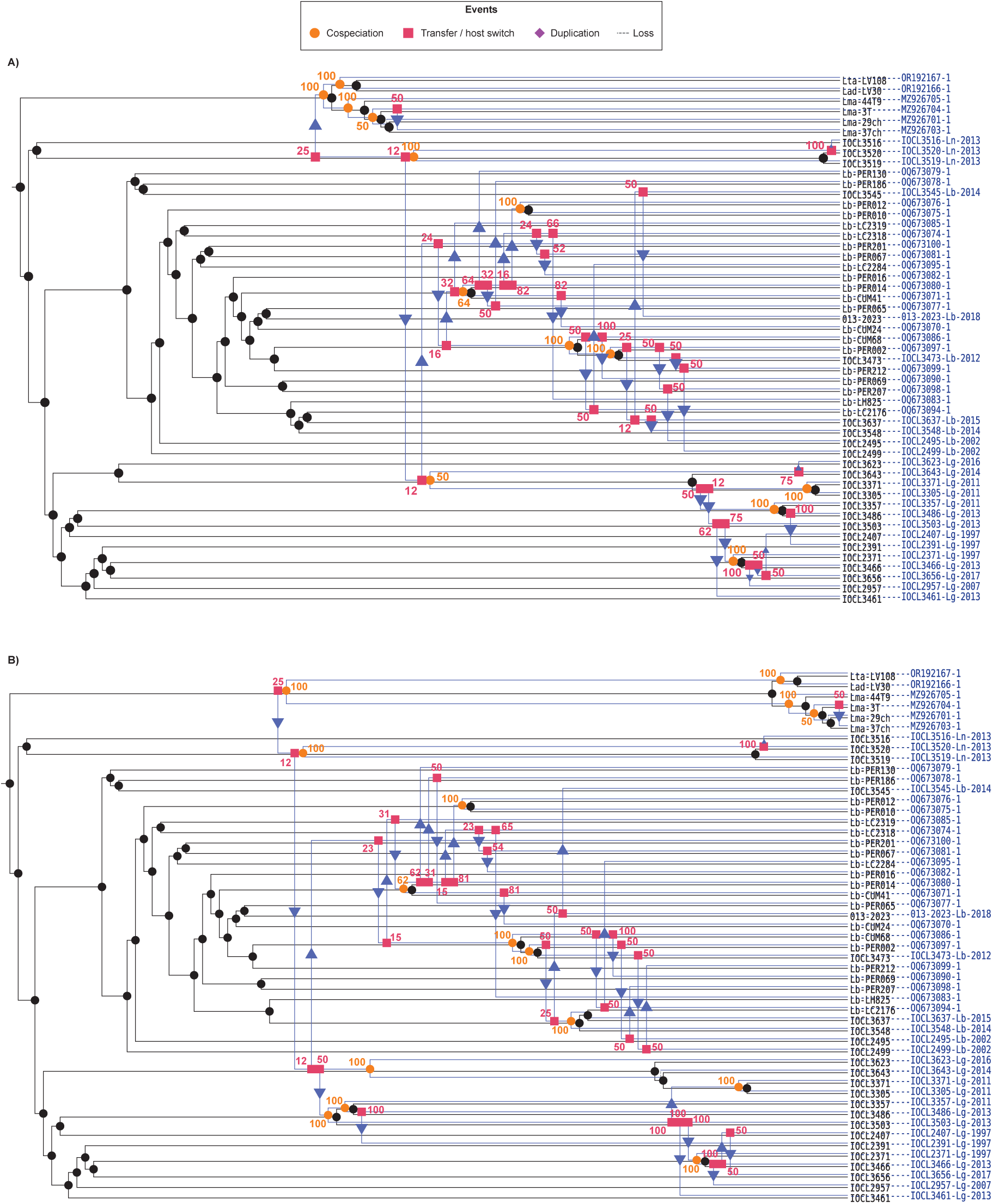
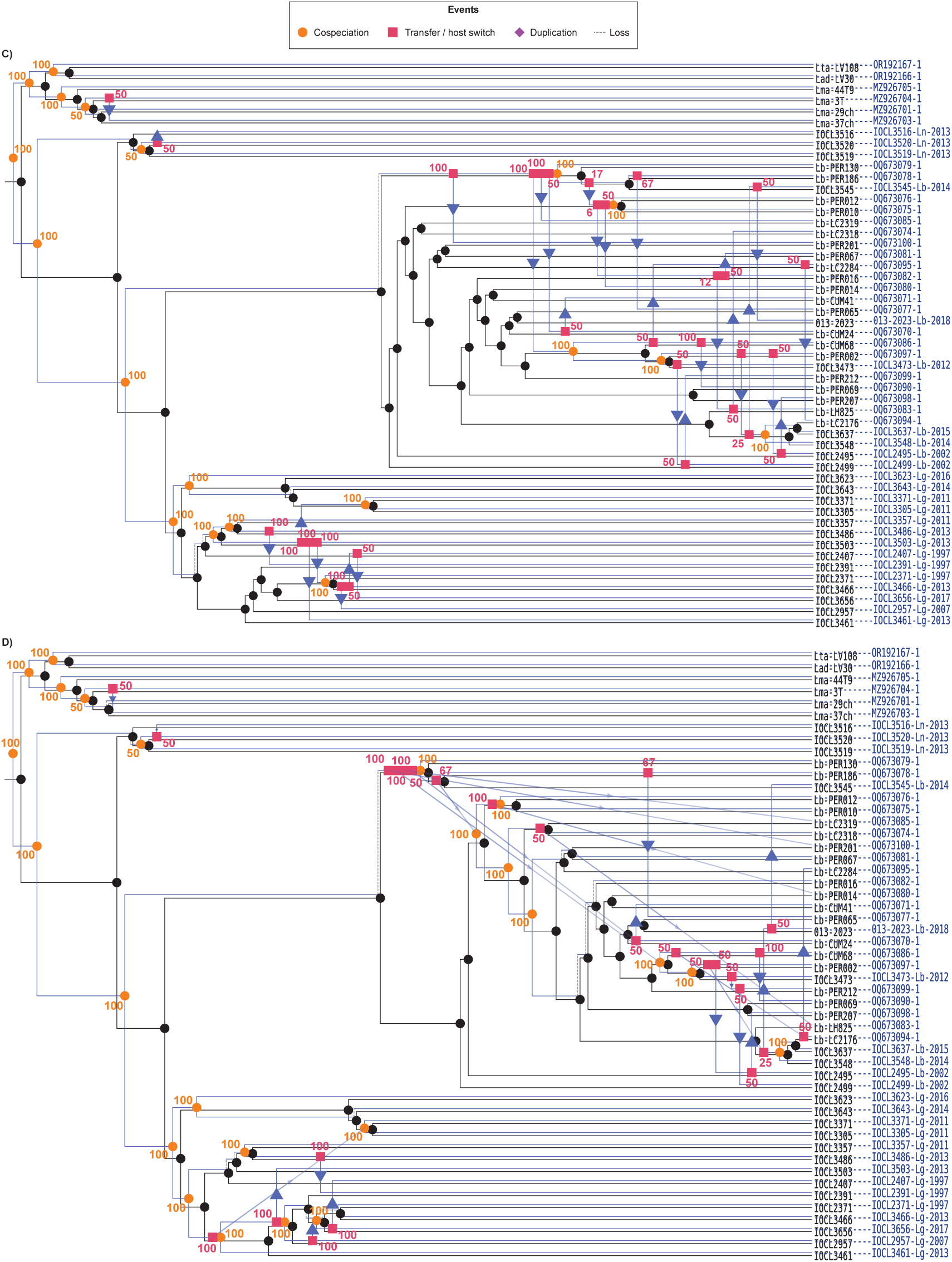
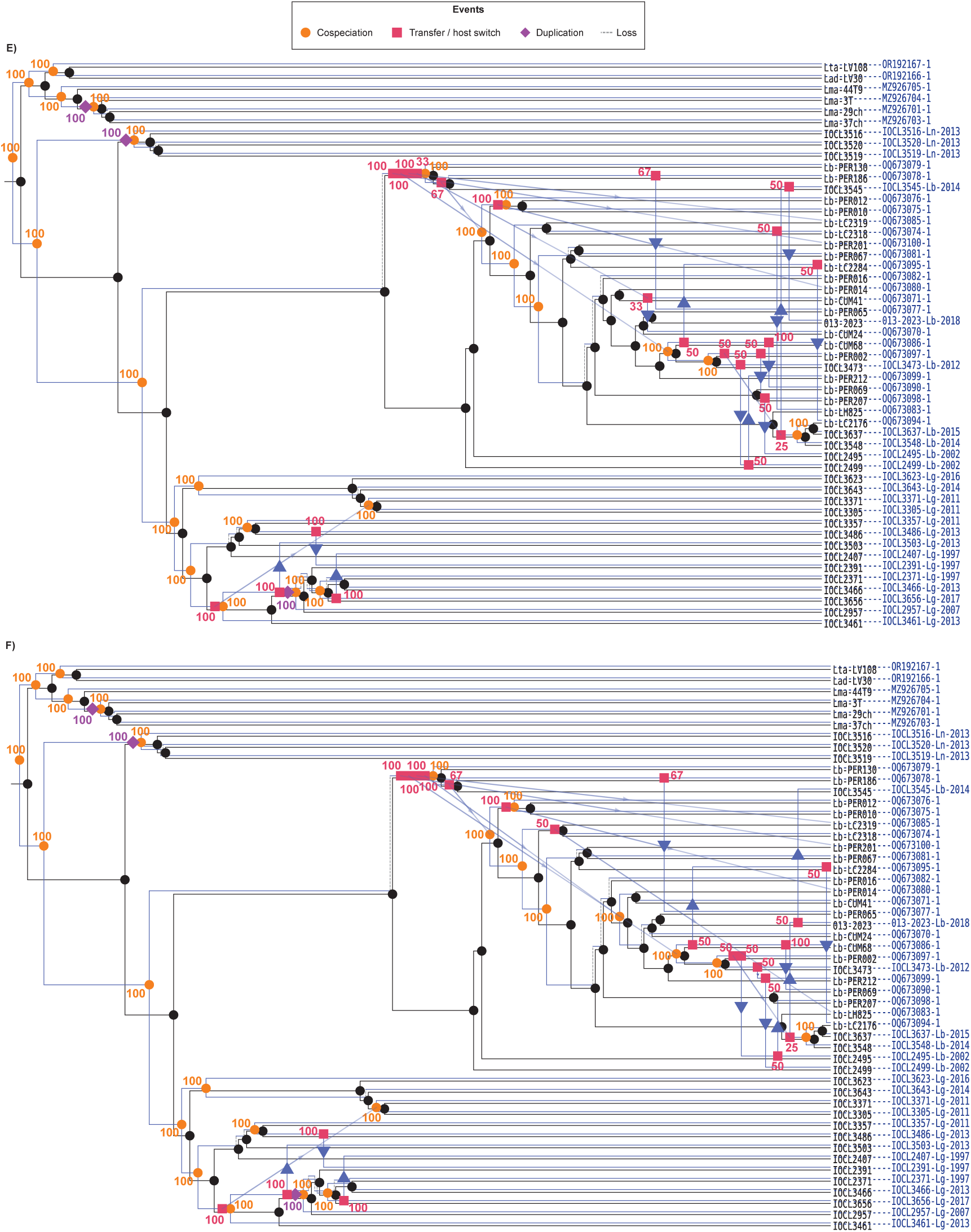
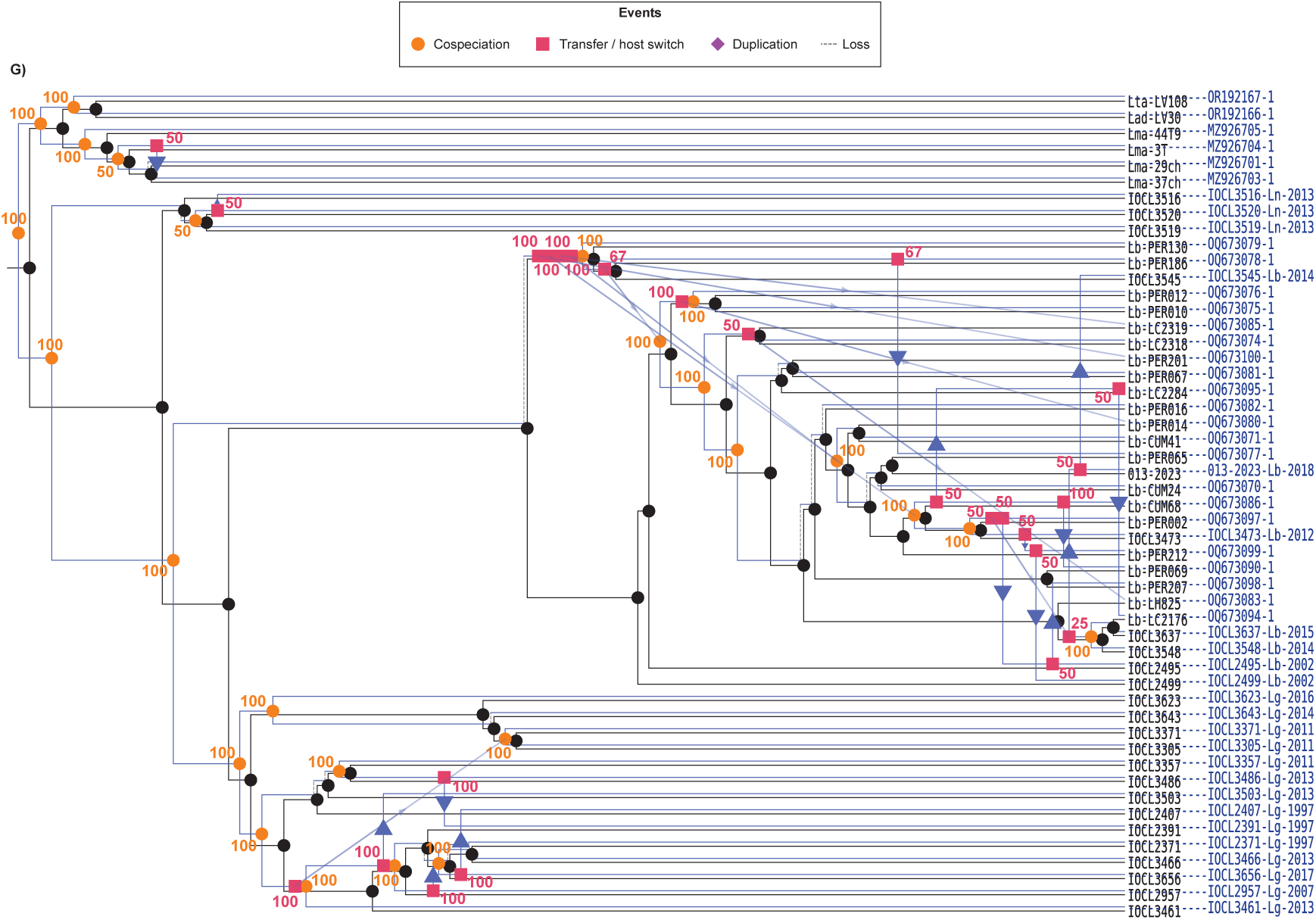
Event-based co-phylogenies of the *Leishmania* divergence, across different subgenera (*L.* (*Viannia*), *L.* (*Leishmania*), *L.* (*Sauroleishmania*)), and LRV species (LRV1, LRV2). Each reconciliation was calculated with different sets of event-costs (S. Fig. 3a; S. Tab. 11): **A)** set 1, **B)** set 2. Support values for the inferred events were based on 100 bootstrap replicates. Panels **C-G** are depicted on the following pages.

**S. Fig. 6.**
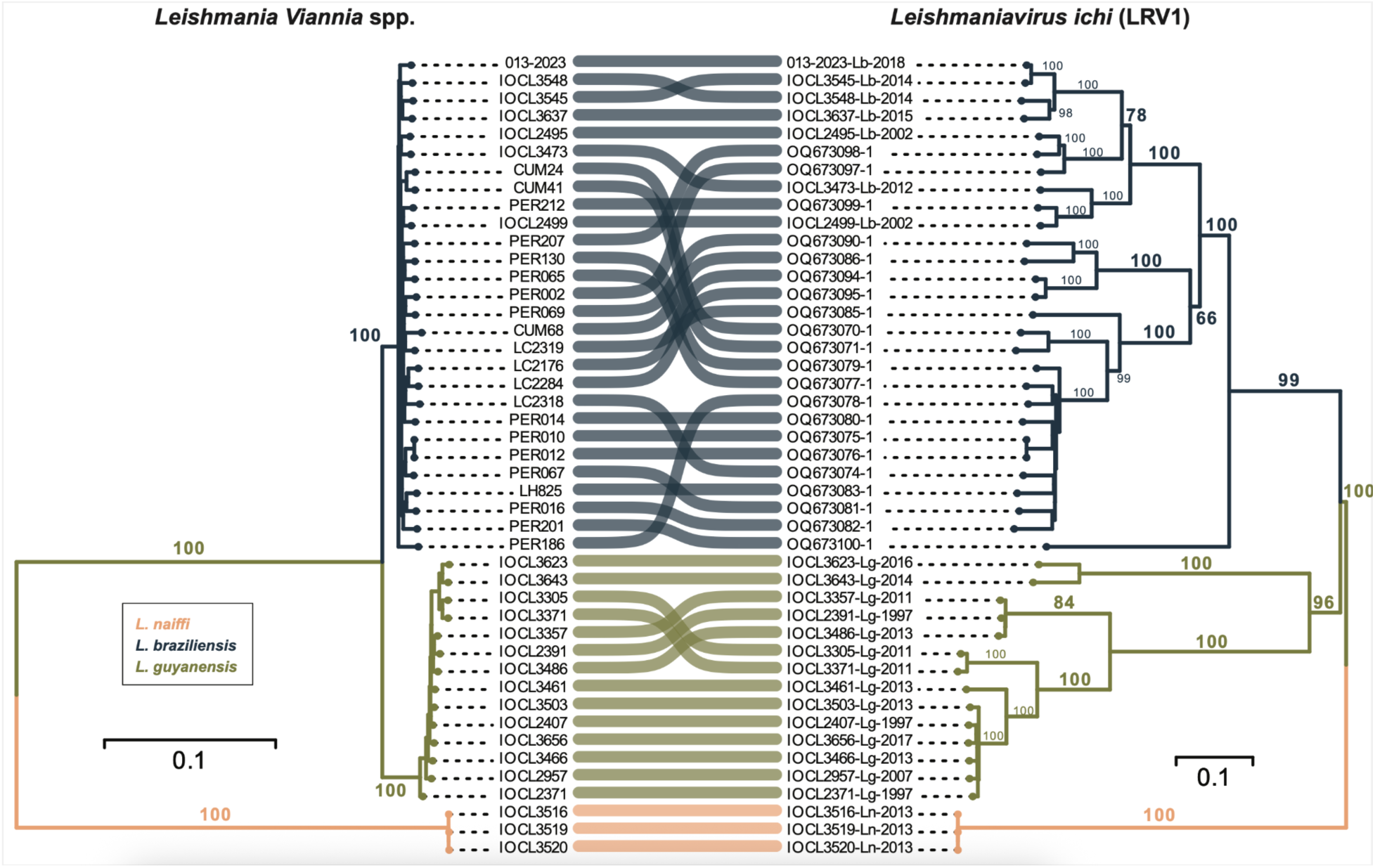
Tanglegram of the *Leishmania Viannia* subgenus and LRV1 phylogenies based on 45 LRV1-positive *L. Viannia* isolates. The *Leishmania* phylogeny was inferred from an alignment of 97,530 concatenated SNPs, corrected for ascertainment bias. The LRV1 phylogeny was inferred from a concatenation of the two protein-coding gene sequences, CP and RDRP (ORF2 and 3, respectively). Colors on the tree are set for visual purposes and do not serve any information on the ancestral states (i.e. internal nodes). Branch values represent bootstrap support values from 10,000 ultra-fast approximated bootstrap replicates. The scale bar depicts the number of substitutions per site.

**S. Fig. 7.**
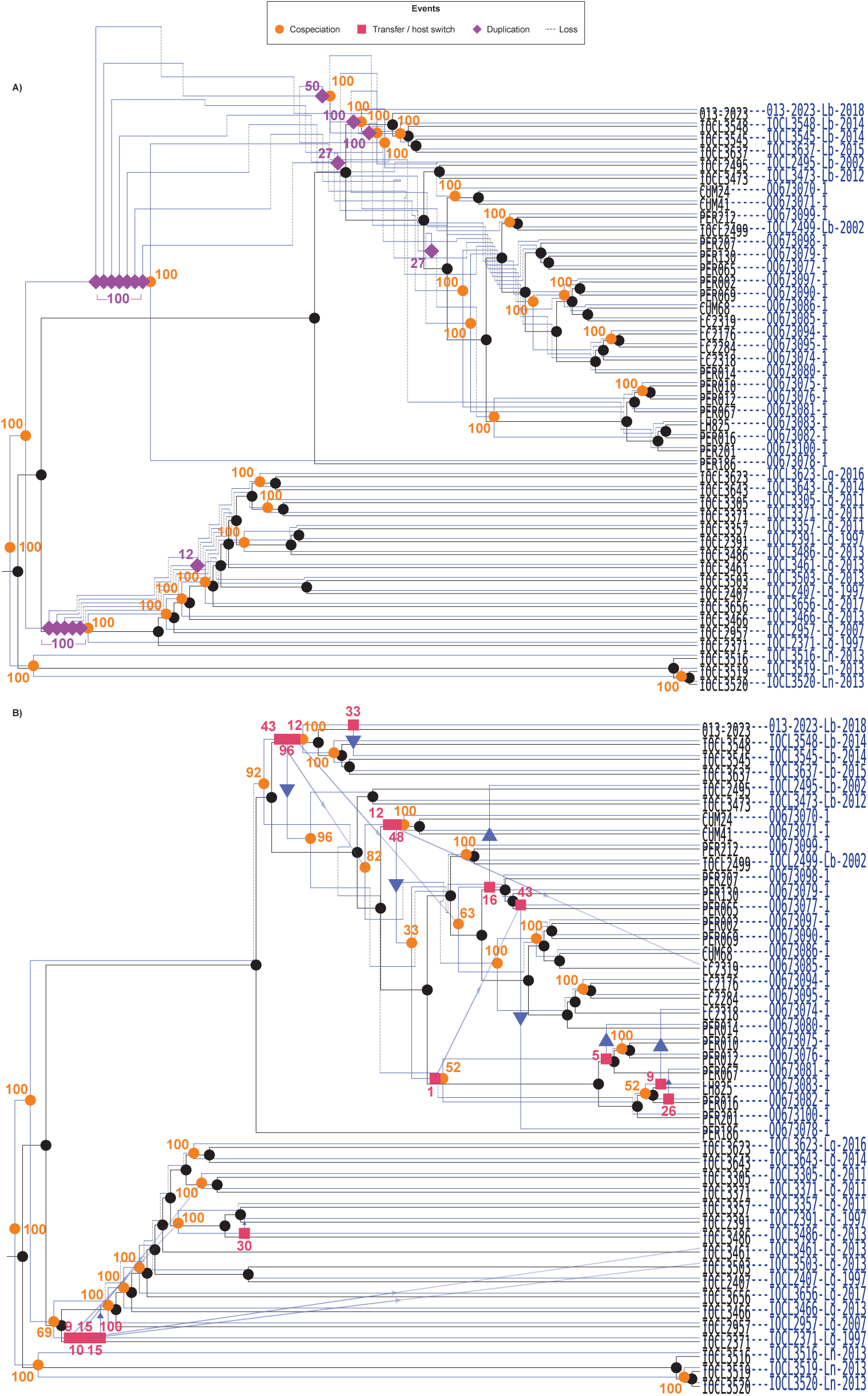
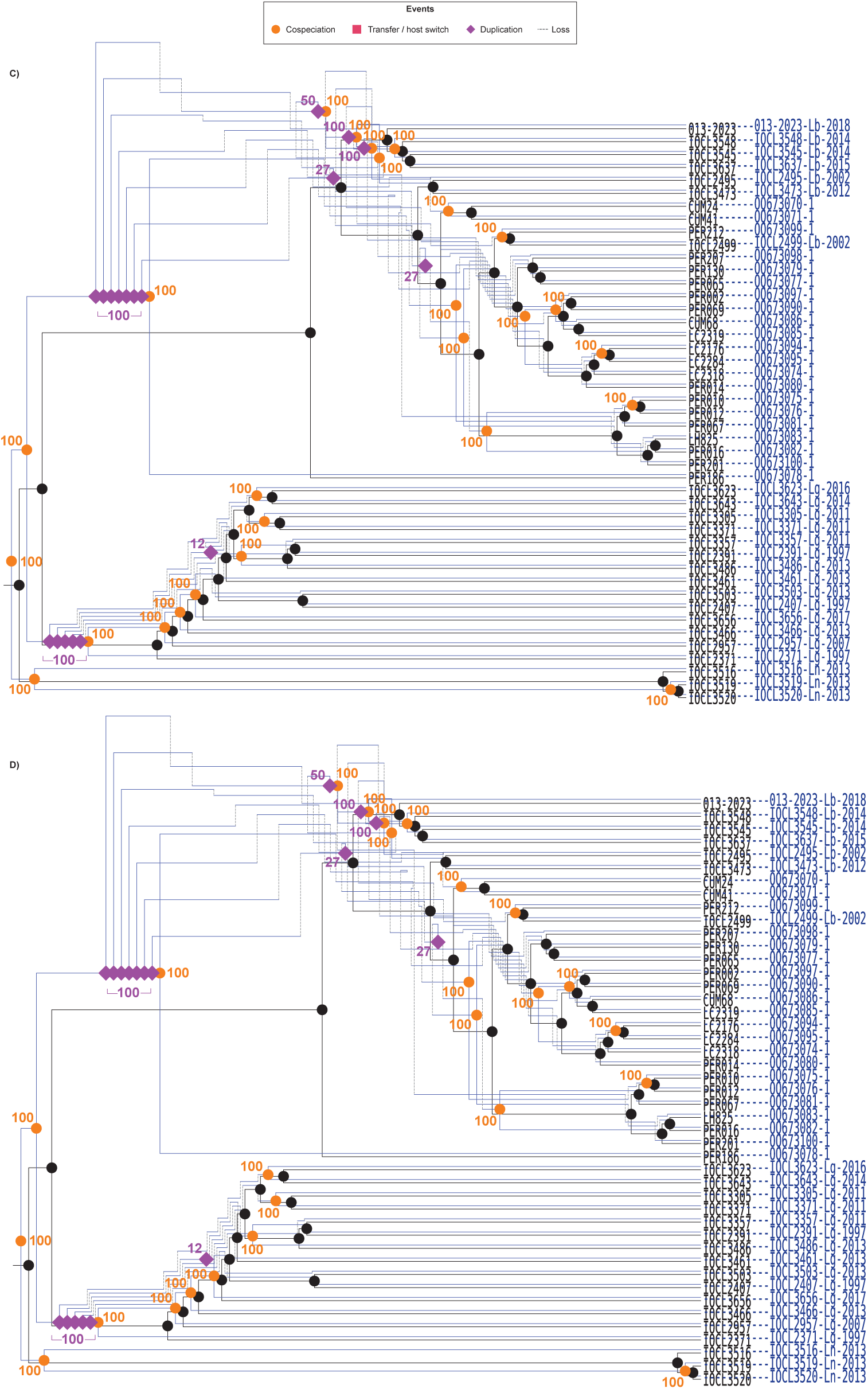
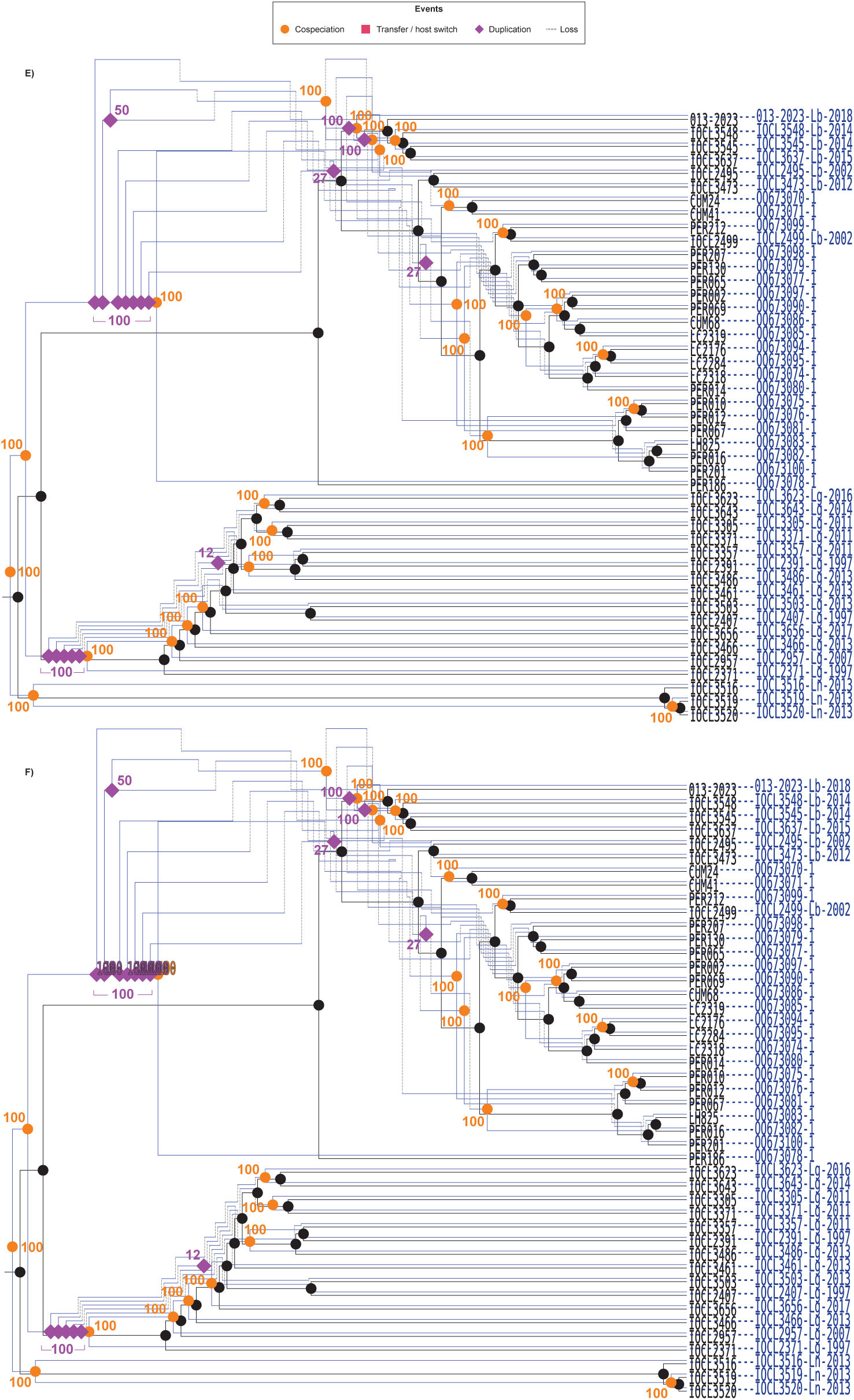
Event-based co-phylogenies of *Leishmania Viannia* and LRV1. Each reconciliation was calculated using different sets of event-costs (S. Fig. 3b;S. Tab. 12): **A)** set 1, **B)** set 2. Support values for the inferred events were estimated from 100 bootstrap replicates. Panels **C-F** are depicted on the following pages.

**S. Fig. 8.**
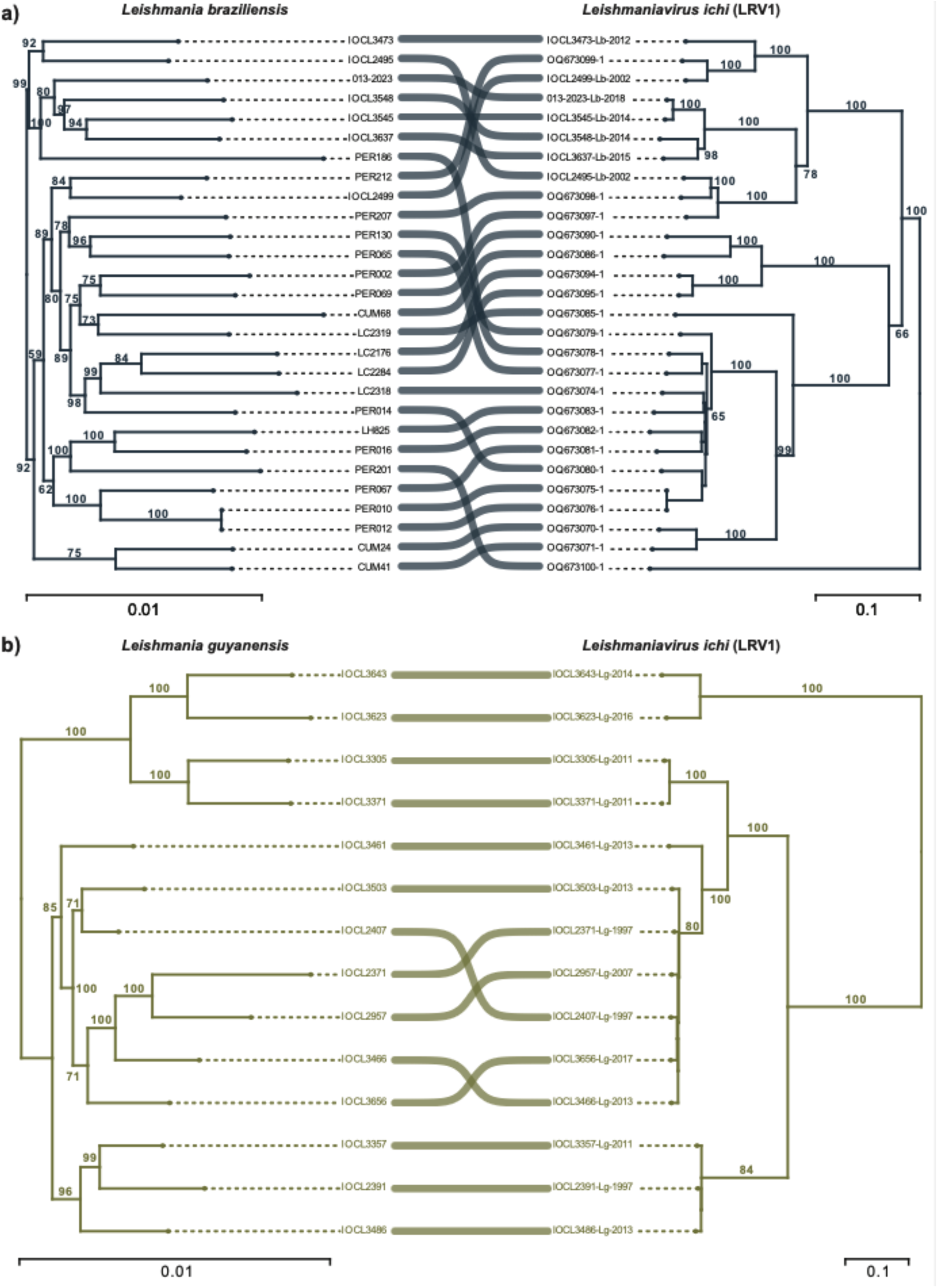
Species-specific tanglegrams of *Leishmania* (*V.*) *braziliensis* (**a**) and *L.* (*V.*) *guyanensis* (**b**) and their associated LRV1 lineages. The *Leishmania* phylogenies represent species-specific partitions of the *L.* (*Viannia*) SNP-based phylogeny (Suppl. Figure 5). Similarly, the LRV1 phylogenies represent specific partitions of the LRV1 phylogeny (Suppl. Figure 5). Branch values represent bootstrap support values from 10,000 ultra-fast approximated bootstrap replicates. The scale bar depicts the number of substitutions per site.

**S. Fig. 9.**
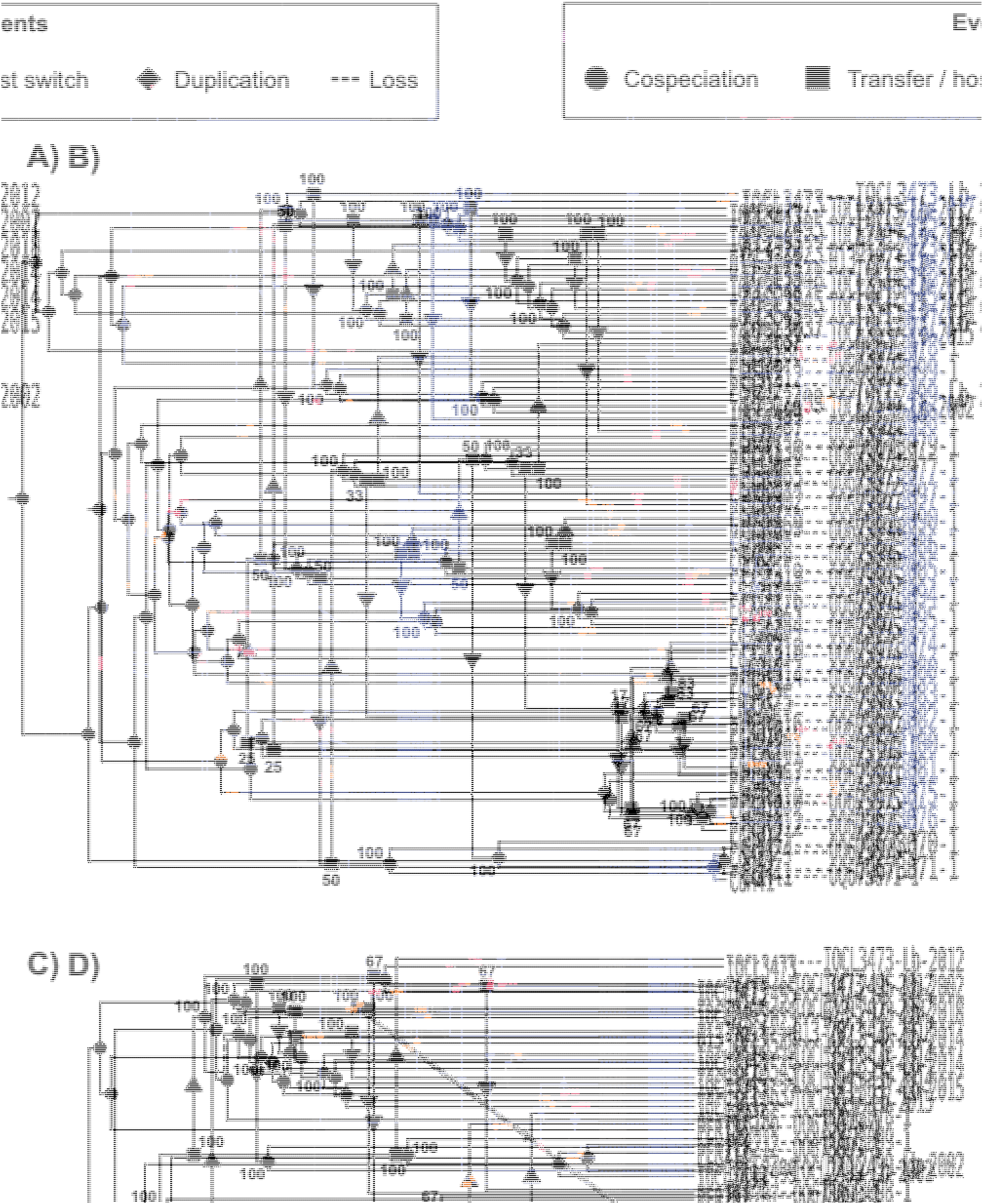
Event-based co-phylogenies of *Leishmania* (*V.*) *braziliensis* and LRV1. Each reconciliation was calculated from different sets of event-costs (S. Fig. 3c; S. Tab. 13): **A)** set 1, **B)** set 2, **C)** set 3, **D)** set 4, **E)** set 5. Support values for the inferred events were estimated from 100 bootstrap replicates.

**S. Fig. 10.**
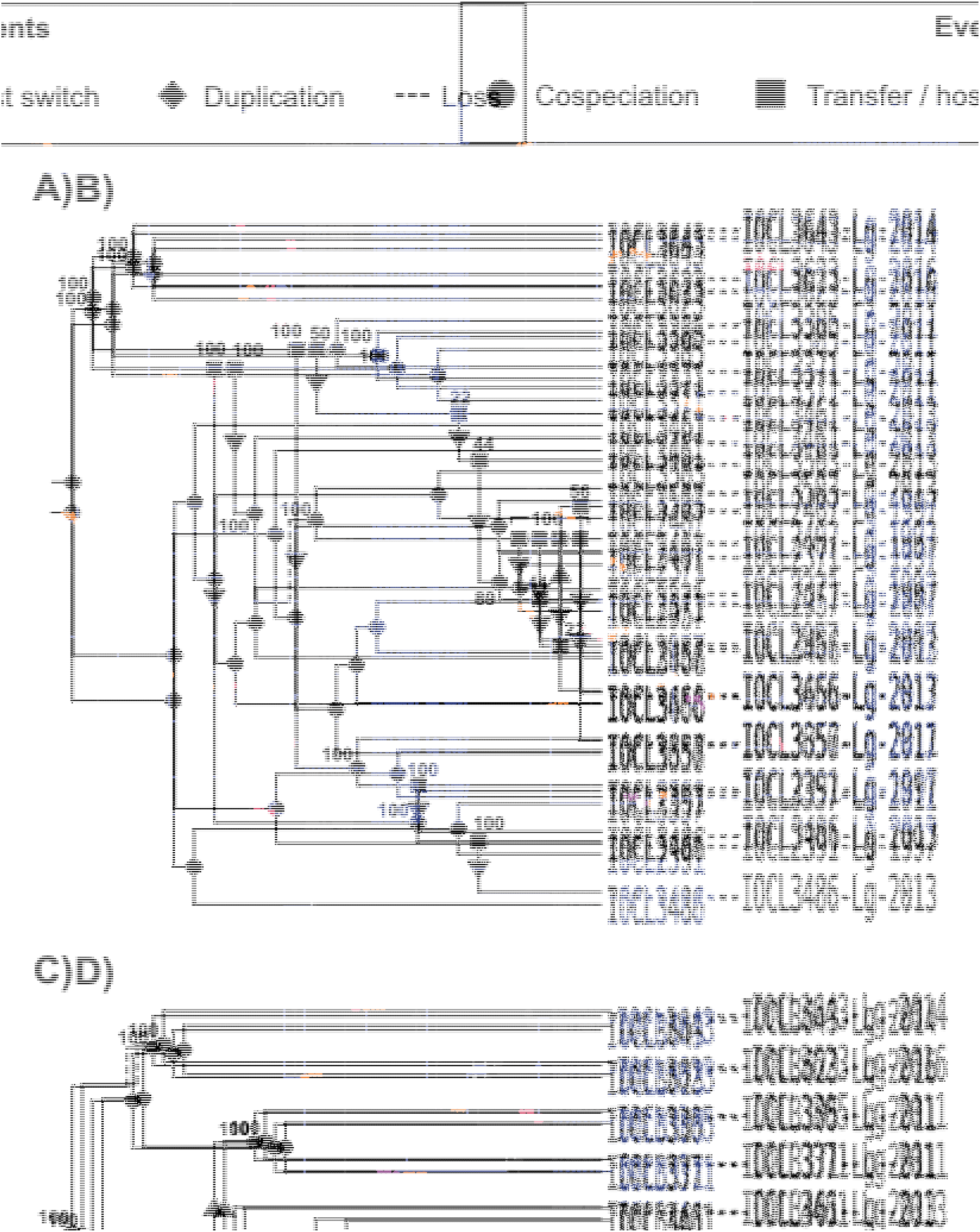
Event-based co-phylogenies of *Leishmania* (*V.*) *guyanensis* and LRV1. Each reconciliation was calculated using different sets of event-costs (S. Fig. 3d; S. Tab. 13): **A)** set 1, **B)** set 2, **C)** set 3, **D)** set 4, **E)** set 5. Support values for the inferred events were estimated from 100 bootstrap replicates. **Supplemental Tables** See excel file.

## Notes

### Competing Interest Statement

The authors have declared no competing interest.

### Summary of Updates

Based on the peer review process regarding the evaluation of the manuscript the manuscript underwent a substantial revision. The main changes were made within the discussion section where a more in depth reflection is made on the co-phylogenetic inferences, including more attention towards its limitations. Secondly, a more nuanced interpretation was made regarding the findings; i.e., less bold statements were used. This new version of the manuscript has a better and more nuanced balance, fitting better with the results and the data that was analysed. Additionally, information on the sequencing data and where data was deposited was added in this new version. Data will be made available upon publication of the manuscript.

